# Confirmation and Transcriptomic Characterization of Glufosinate-ammonium Resistance in Waterhemp (*Amaranthus tuberculatus*) Populations from Illinois

**DOI:** 10.64898/2026.02.06.704380

**Authors:** Isabel Werle Noe, Cristiana Bernardi Rankrape, Logan Miller, Eduardo Lago, Rishabh Singh, Alexander J. Lopez, Aaron G. Hager, Karla L. Gage, Patrick J. Tranel

**Author notes:** **Author for correspondence:** Patrick J. Tranel, University of Illinois Urbana-Champaign, 1201 West Gregory Drive, Urbana, IL 61801. These authors contributed equally to this work (Isabel Werle Noe, Cristiana Bernardi Rankrape, Logan Miller).

## Abstract

Glufosinate-ammonium (GA) has been widely used in Midwestern fields, and in recent years a growing number of failures to control waterhemp [*Amaranthus tuberculatus* (Moq.) Sauer] have raised concerns about the potential evolution of resistance. The goal of this study was to investigate four independent cases of suspected resistance to GA in *A. tuberculatus* from Illinois using greenhouse, field, and transcriptomics studies. Greenhouse dose-response experiments revealed resistance ratios ranging from 2.2- to 3.4-fold based on survival and from 1.3- to 2.8-fold based on dry biomass relative to a susceptible population. A subsequent field study where one of the populations originated confirmed that twenty percent of treated plants survived the labeled GA field-recommended rate. Screening for other herbicide sites of action revealed that most populations showed reduced sensitivity to atrazine, glyphosate, and imazethapyr, surviving up to three times the field-recommended rates, and to a lesser extent, lactofen and fomesafen. Transcriptomic analysis of plants surviving GA revealed no resistance-associated mutations or differential transcript abundance in the plastidic and cytosolic isoforms of glutamine synthetase. Among the four suspected resistant populations, there were 182 genes differentially expressed relative to two susceptible populations. Different sets of genes were differentially expressed among the populations studied, with only one gene (upregulated relative to two susceptible populations) shared among all four. Many of the differentially expressed genes, including cytochrome P450s, glutathione *S*-transferases, glycosyltransferases, transporters, and transcriptional regulators, are commonly associated with metabolic resistance. Gene ontology enrichment analyses indicated significant overrepresentation of stress response, defense regulation, and secondary metabolism categories across the populations. Together, these findings provide evidence for the evolution of GA resistance in populations of *A. tuberculatus* in Illinois. While more in-depth studies are needed to fully characterize the underlying mechanisms, the consistent differential expression of metabolism-related genes and no indication of target-site mechanisms points to a potential metabolic basis for resistance.

## Introduction

The use of glufosinate-ammonium (GA) has increased substantially in the United States over the past decade. Between 2010 and 2018, following the introduction of GA-resistant soybean varieties, total use of GA-based products increased from less than 2.3 million kg ai in 2010 to more than 4.5 million kg ai after 2016 (USGS 2018), of which most GA applications have been concentrated in the Midwest and Southern regions (Baker and Stone 2015), where soybean and corn production are more prominent. Inconsistent weed control with GA has been frequently reported in recent years, often raising concern whether this variability is solely due to environmental conditions and other factors such as weed size at application, or if it signals the early evolution of GA resistance in weed populations.

Glufosinate-ammonium is a non-selective, post-emergence herbicide with rapid activity leading to the onset of visible symptoms within hours of application. The primary mode of action of GA is the inhibition of glutamine synthetase (GS, EC 6.3.1.2). The inhibition of GS following GA application leads to ammonia accumulation, followed by a feedback inhibition of photorespiration, which results in the generation of reactive oxygen species (ROS), particularly hydrogen peroxide. The resulting oxidative stress triggers lipid peroxidation, which affects membrane integrity and ultimately causes cell death (Takano et al. 2019, 2020a). Despite its rapid activity, some physicochemical properties of GA can substantially affect its performance. GA is a highly hydrophilic compound and has a low octanol-water partition coefficient (log Kow ≈ −4.5; log P: −5.0), which limits its ability to passively enter lipid membranes and restrict systemic movement (Brown and Farenhorst 2024). Unlike glyphosate, which despite similar hydrophilic properties (log Kow: −3.5; log P: −4.6) is efficiently transported via the phloem through phosphate transporters, GA lacks an efficient transmembrane transporter (Brown and Farenhorst 2024; Okada et al. 2019; Takano et al. 2020b). As a result, GA is primarily translocated through the xylem via the apoplast and driven by transpiration streams (Takano et al. 2020b).

Because of limited translocation, thorough foliar coverage is essential for adequate weed control with GA. However, even with adequate coverage, environmental conditions such as light intensity, relative humidity, and temperature can greatly influence the efficacy of GA in the field. Light is essential for hydrogen peroxide production in photosynthetic light reaction centers, which is critical for GA-induced phytotoxicity and GS expression and abundance (Takano and Dayan 2021). A recent study by Landau et al. (2025) using random forest models trained on over ten thousand field trials demonstrated that under low solar radiation conditions, the probability of successful control of waterhemp [*Amaranthus tuberculatus* (Moq.) Sauer], morningglory species (*Ipomoea* spp.), and giant foxtail (*Setaria faberi* Herrm.) was negatively impacted. Relative humidity can affect herbicide absorption, especially shortly before or after GA application (Landau et al. 2025; Ramsey et al. 2006). Under elevated relative humidity conditions, the drying time of GA droplets is extended, which increases foliar uptake, and the cuticle is more hydrated, leading to greater absorption of hydrophilic compounds such as GA (Coetzer et al. 2001). Temperature is another important factor pertaining to its effect on plant metabolic processes and herbicide translocation. Typically, studies point to an increased basipetal translocation of GA at warmer temperatures while most of the herbicide tends to be translocated to the tip of the leaves at cooler temperatures (Kumaratilake and Preston 2005). Current recommendations emphasize applying GA under high temperature and high humidity conditions to maximize efficacy (Lingenfelter 2024; Preston 2024; Singh et al. 2024).

In addition to environmental factors, morphological aspects of different weed species and weed size at the time of application can influence their sensitivity to GA (Steckel et al. 1997). The label-recommended size for GA to effectively control *Amaranthus* species is 10 cm or less, with larger plants (>15 cm) requiring higher rates (Anonymous 2024) and, often, sequential applications are required to achieve desirable control. The type of weed (e.g., broadleaf vs grass species) also plays a role in GA efficacy (Haarmann et al. 2020; Meyer and Norsworthy 2020). For instance, dicotyledonous species are generally more susceptible to GA than monocots due to differences in leaf morphology and cuticle composition (Takano and Dayan 2020). Furthermore, some weed biotypes may reduce GA efficacy through variation in architectural traits, such as the rate of axillary bud development. These buds, often located near the plant base, may escape herbicide contact and facilitate regrowth after treatment, which is frequently observed following field applications of GA.

To date, resistance to GA has been confirmed in six weed species (Heap 2025). Among these, Palmer amaranth (*Amaranthus palmeri* S. Watson) is the only dicot confirmed to have evolved resistance to GA (Jones et al. 2024; Noguera et al. 2022; Priess et al. 2022). In two of the three documented GA-resistant *A. palmeri* cases, resistance was conferred by overexpression of the GS target gene through an interesting mechanism involving extrachromosomal circular DNA (eccDNA) (Carvalho-Moore et al. 2025, 2022; Noguera et al. 2024, 2022). While resistance to GA in *A. tuberculatus* has not yet been confirmed, increasing reports of reduced GA efficacy in Illinois populations warrant investigation. In this study, we evaluated *A. tuberculatus* populations with reported reduced sensitivity to GA from four Illinois counties using greenhouse and field experiments to investigate potential resistance to GA. Furthermore, we evaluated control of these populations with alternative herbicide sites-of-action (SOA) in additional greenhouse experiments to identify suitable alternatives. We also assessed changes in the coding sequence of GS genes and constitutive variation in transcriptome profiles to gain initial insights into potential mechanisms underlying reduced sensitivity to GA in these populations.

## Materials and Methods

### Plant Materials

Seeds were collected from fields reporting failure to control *A. tuberculatus* with labeled rates of GA in Carroll (CAR), Kankakee (SDY), McLean (M01), and Franklin (FRA) counties in Illinois between 2022 and 2024 (Figure 1). Two susceptible populations, WUS (Brown County, Ohio; GRIN ID: PI 698378) and BRC (SIU Belleville Research Center, St. Clair County, Illinois), were included in the transcriptomics study, and only WUS was included in the greenhouse dose-response studies. To reduce heterogeneity of the suspected resistant populations, we crossed individuals from each suspected resistant population that had survived GA at the field-recommended rate (654 g ai ha⁻¹; Liberty® 280 SL herbicide, BASF Corporation) under greenhouse conditions. For each population, one male and at least three female survivors were placed in an enclosed environment to allow for cross-pollination. Seeds were collected separately from each female plant and screened a second time with GA at the same rate. Seeds derived from females with the most homogeneous survival response in the second screening were pooled to form uniform resistant populations for use in all subsequent greenhouse experiments.

**Figure 1.**
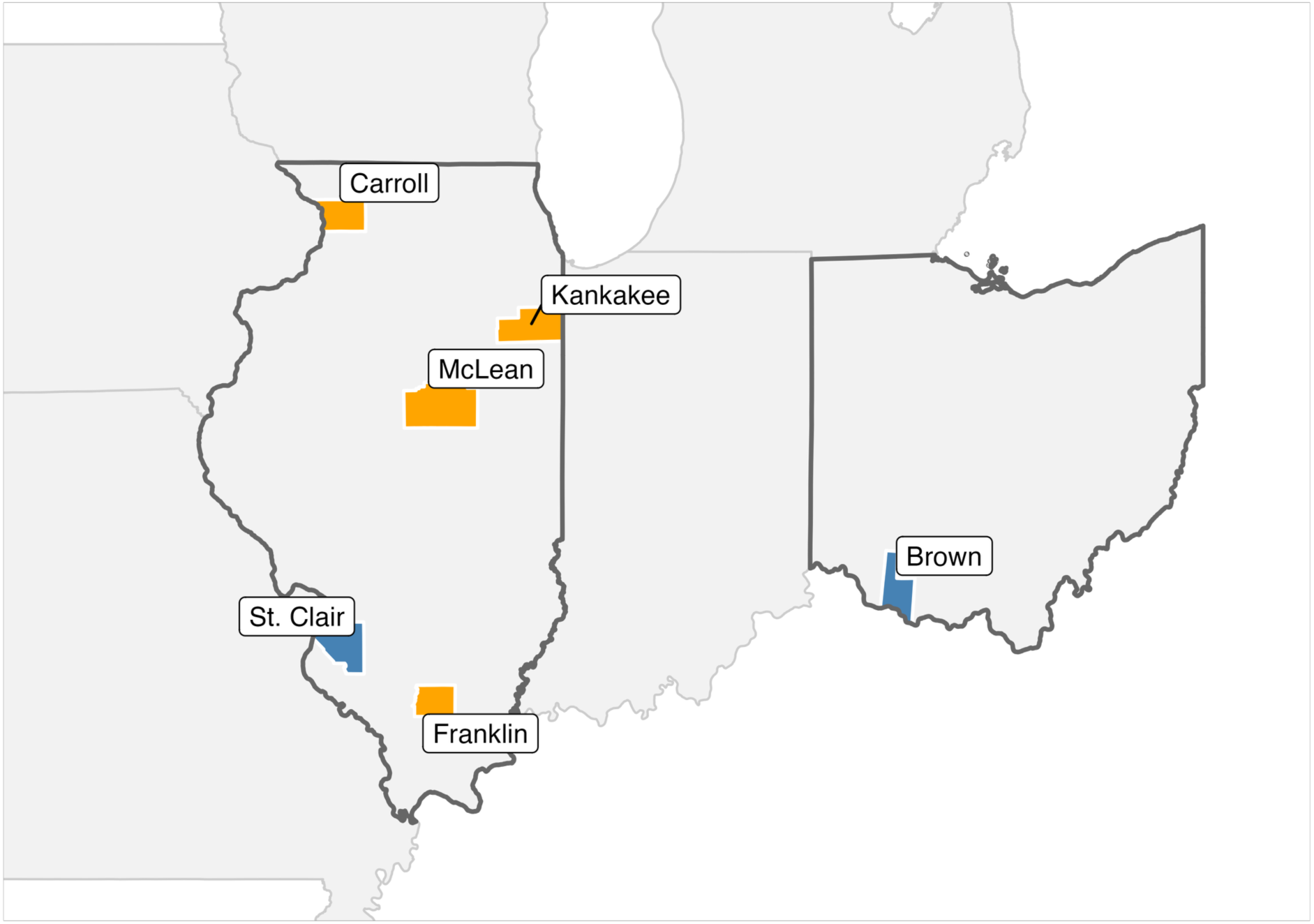
Map showing the geographic distribution of *Amaranthus tuberculatus* populations used in this study. Suspected-resistant populations were collected from Carroll (CAR), Kankakee (SDY), McLean (M01), and Franklin (FRA) counties in Illinois. The susceptible populations BRC and WUS were collected from St. Clair County, Illinois, and Brown County, Ohio, respectively. Counties where suspected resistant populations originated are shown in orange and susceptible populations in blue.

### Greenhouse Conditions

Greenhouse studies for the CAR, SDY, and M01 populations were conducted at the University of Illinois Plant Care Facility (PCF), while those for the FRA population were carried out at the Southern Illinois University Horticultural Research Center (HRC), with slight variations in greenhouse conditions, soil type, and fertilization regimes between the two sites. For all experiments described herein, at the PCF, greenhouse conditions were maintained under a 16-hour photoperiod, with supplemental LED lighting (extended white, 1000 µmol m^-2^ s^-1^) used to extend day length and automatically deactivated when outside light intensity exceeded 600 W m⁻². Daytime temperatures ranged from 28 to 30 °C and nighttime temperatures from 24 to 26 °C, and relative humidity ranged from 30 to 63% during the day and from 38 to 64% during the night. A custom sandy loam potting mix containing 70% sand, 15% silt, 15% clay, and 7% organic matter with a pH of 6.3 was used for all experiments at the PCF. At the HRC site, greenhouse settings were maintained at a 16-hour photoperiod, and light was supplemented with 1000W High Pressure Sodium (HPS) fixtures. The daytime temperatures ranged from 25 to 40°C, and nighttime temperatures from 20 to 27°C. The daytime relative humidity ranged from 37 to 74% and nighttime relative humidity from 60 to 88%. A growing medium potting mix containing approximately 75–85% peat moss organic matter, and 10–20% perlite and a pH range of 4.5 to 6.0 was used for all experiments at the HRC.

### Greenhouse Dose-Response Studies

Whole-plant dose-response assays were conducted to evaluate the responses of the suspected resistant populations CAR, SDY, M01, and FRA to GA. The susceptible population WUS was included for comparison in dose-response studies at both locations. Unless otherwise noted, the experimental procedures described herein were consistent across both facilities. Seeds from each population were sown into 164 cm³ containers containing a custom potting mix and maintained in a greenhouse as previously described. The experiments followed a completely randomized design (CRD), with seven replications per treatment, and were repeated once over time. Plants were treated with GA (Liberty® ULTRA Herbicide, BASF Corporation) at 0, 139, 209, 312, 469, 702, 1,054, and 1,581 g ai ha⁻¹. All treatments included ammonium sulfate (AMS) at 2.5% of the application volume.

For greenhouse-based experiments, herbicide treatments were applied using spray chambers equipped with 80015 EVS nozzles (TeeJet® Technologies, Glendale Heights, IL), calibrated to deliver 187 L ha⁻¹ at 207 kPa. Applications were made when *A. tuberculatus* plants reached 7.5 to 10 cm in height. All GA treatments were applied between 12:00 and 1:30 PM.

At 21 days after application (DAA), plant survival was recorded as 0 (dead) or 1 (alive). Above ground biomass was harvested separately for each plant, oven-dried at 40 °C for five days at PCF and at 60 °C for three days at HRC and then weighed.

### Field Study

A field dose-response study was conducted at the original collection site of the CAR population in Carroll County, Illinois. The experiment followed a randomized complete block design (RCBD) with three replications per treatment and was conducted during a single growing season in 2025. Individual plots measured 3 m by 9 m. The field location was planted with corn, and the entire field received a preemergence application of isoxaflutole + flufenacet + thiencarbazone-methyl (511 g ai ha⁻¹; Trivolt®, Bayer Crop Science) and atrazine (1,120 g ai ha⁻¹; Infantry™, Growmark Inc.).

On the day of the GA dose-response application, corn plants in the experimental area were flattened manually, and 10 to 15 *A. tuberculatus* plants between 7.5 and 12.5 cm in height were randomly selected and marked with a stake in each experimental plot. Treatments applied to the CAR population included 0, 186, 371, 742, 1,484, 2,968, and 5,936 g ai ha^−1^ of GA (Liberty® ULTRA Herbicide). All treatments included AMS at 2.5% v/v. Herbicide applications were made using a CO_2_ pressurized backpack sprayer equipped with TT 110025 TeeJet® nozzles (TeeJet® Technologies, Glendale Heights, IL, USA) spaced 50 cm apart and calibrated to deliver 187 L ha^−1^ at 5.6 km h^−1^ and 248 kPa. At the time of application, air temperature was 31°C, the relative humidity was 71%, and the wind speed was 4.9 m s-^−1^.

To evaluate the response of CAR to GA rates, visual assessments of *A. tuberculatus* control were made on an individual-plant basis for each tagged plant within plots at 6, 12, and 24 DAA, using a scale from 0 (no control) to 100 % (complete control).

### Dose-Response Statistical Analyses

To model the responses of *A. tuberculatus* populations to GA rates, two log-logistic regression models were used. A three-parameter log-logistic model was fitted to plant survival data, and a four-parameter log-logistic model was used for dry biomass data. Both models were implemented using the ‘*drc*’ package (Ritz et al. 2015) in R v4.5.2. The three- and four-parameter log-logistic models followed the respective equations:

Three-parameter model (Equation 1):

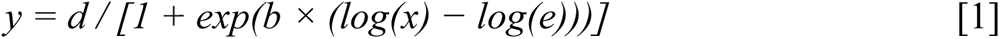

Four-parameter model (Equation 2):

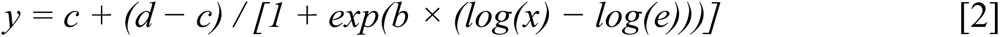

where *y* represents the response variable (survival or dry biomass), *x* is the herbicide rate, *c* is the lower asymptote, *d* is the upper asymptote, *e* is the effective rate that reduces the response by 50% (LD₅₀/GR₅₀), and *b* is the slope around the LD₅₀/GR₅₀.

### Multiple Herbicide Site-of-Action Screening

In addition to dose-response assays, greenhouse experiments were also conducted to assess the responses of each *A. tuberculatus* population to herbicides representing different SOA groups. At the PCF site, CAR, SDY, and M01 were screened with 2,4-D, atrazine, fomesafen, and glyphosate. At the HRC site, FRA and BRC populations were screened with 2,4-D, atrazine, fomesafen, glyphosate, lactofen, imazethapyr and paraquat. Each herbicide was applied separately at 0.5×, 1×, and 3×the field-labeled rate (Supplementary Table S1). Each population and treatment combination had four to six replications, where a replication consisted of an individual plant grown in a 70 x 70 x 90 mm (700 cm³) plastic pot. Experiments were arranged in a CRD. Screenings with the FRA population were conducted twice, and CAR, SDY, and M01 were tested in a single experimental run due to limited seed germination and availability. Detailed herbicide information and rates are provided in Supplementary Table S1.

### Whole Transcriptome Sequencing

To investigate variation in constitutive gene expression among populations of *A. tuberculatus,* an RNA sequencing (RNA-Seq) study was conducted. Plants from field-collected seeds of the suspected resistant populations CAR, SDY, FRA, and M01, and the two herbicide-sensitive populations, WUS and BRC, were grown under previously described greenhouse conditions at the PCF. Tissue samples were collected from a single, fully developed leaf from each of four biological replicates of each population prior to GA treatment. Leaf samples were frozen in liquid nitrogen upon collection and stored at −80°C until RNA extraction. After tissue collection, plants were treated with the labeled rate of GA (654 g ai ha⁻¹; Liberty® 280 SL Herbicide). At 21 DAA, plants were phenotyped for survival and classified as 0 (dead) or 1 (alive). Only samples from plants of the suspected resistant populations surviving GA, as well as those from susceptible plants that were completely controlled were retained for the RNA-Seq study.

Total RNA was extracted using a TRIzol-based protocol, and each sample was treated with DNase I to remove genomic DNA contamination (Simms et al. 1993). Overall RNA quality was checked on a 1% agarose gel, and RNA concentrations were measured using a Qubit v4.0 Fluorometer (Thermo Fisher Scientific, Waltham, MA, USA). Libraries were sent to the Roy J. Carver Biotechnology Center, University of Illinois and sequenced on an Illumina NovaSeq X Plus 10B platform, producing 150-bp paired-end reads. Sequencing data have been deposited in the NCBI Sequence Read Archive (SRA) under BioProject ID PRJNA1348745 (BioSamples SAMN2917128-SAMN2917151).

### Sequencing Data Processing and Differential Gene Expression Analysis

Read quality was assessed with FastQC v0.12.0 (https://www.bioinformatics.babraham.ac.uk/projects/fastqc/) and summarized using MultiQC v1.12 (Ewels et al. 2016). Reads were aligned to an available *A. tuberculatus* reference genome (Raiyemo et al. 2024) using STAR v2.7.10b (Dobin et al. 2013), and transcript quantification was performed with featureCounts v2.0.6 (Liao et al. 2014) from the Subread package. Raw transcript counts were converted to counts per million (CPM) and transcripts with fewer than one CPM in at least two samples per population were considered lowly expressed and removed from further analysis. Differences in sequencing depth between samples were accounted for using the Trimmed Mean of M-values (TMM) normalization. Factor analysis on control samples was conducted for normalization of read counts using the unwanted variation (RUV) method using RUVSeq v1.36.0 (Risso et al. 2014).

Differential gene expression (DGE) analysis was conducted using edgeR v4.0.16 (Robinson et al. 2010), applying the quasi-likelihood framework (QLF) within a generalized linear model (GLM). Differences in constitutive gene expression were initially tested by separately comparing each suspected-resistant *A. tuberculatus* population against each of the susceptible populations, WUS and BRC. Genes retained for downstream analyses were required to meet three criteria: (1) significance at a 5% false discovery rate (FDR), (2) a minimum |log fold| change of 1, and (3) consistent differential expression in comparisons against both susceptible populations.

### Functional Annotation of Differentially Expressed Genes and Heat Map Analysis

Selected differentially expressed genes were used for gene ontology (GO) functional enrichment and heat map construction. Protein sequences of *A. tuberculatus* genes were submitted to eggNOG-mapper v2.1.12 (Cantalapiedra et al. 2021) for ontology annotations, and GO enrichment was conducted using the R package topGO v2.54.0 (Alexa and Rahnenführer 2009) using the classic Fisher’s exact test across biological process (BP), cellular component (CC), and molecular function (MF) categories. The list of genes consisted of all annotated genes with GO terms, and a node size filter of 10 was applied for GO term selection. *P*-values were corrected using the Benjamini-Hochberg procedure, and significantly enriched GO terms were recorded for each population. For heat map construction, hierarchical clustering was performed using correlation-based distance metrics using pheatmap v1.0.13 (Kolde and Kolde 2015).

### Identification of Glutamine Synthetase Homologs and Resistance Mutations

To identify mutations that may confer target-site resistance to GA in the suspected resistant populations, RNA reads were used to screen for functionally relevant variants within the GS target genes. Genes encoding *GS1* and *GS2* were first identified by querying homologous GS coding sequences obtained from NCBI against the *A. tuberculatus* reference genome using BLASTn. Homologous GS genes were subsequently identified in *A. hybridus* (Raiyemo et al. 2025), *A. hypochondriacus* (Lightfoot et al. 2017), *A. palmeri* (Raiyemo et al. 2025), *A.* retroflexus (Raiyemo et al. 2025), and *A. tricolor* (Wang et al. 2023). Protein sequences from the six *Amaranthus* species, together with sequences from *Chenopodium quinoa*, *Sonchus oleraceus*, *Setaria italica*, and *Oryza sativa*, were aligned using MEGA v12.1 (Kumar et al. 2024). Phylogenetic trees were generated and visualized using Interactive Tree of Life (iTOL) v7 (Letunic and Bork 2024). Protein identifiers for genes obtained from NCBI and used in phylogenetic analysis are listed in Supplementary Table S2.

Variant calling was performed on the aligned reads using bcftools v1.18 (Danecek and McCarthy 2017). Variants were normalized against the reference genome and filtered to retain only biallelic single-nucleotide polymorphisms (SNPs) with quality scores greater than 20. Functional effects of SNPs, including synonymous, missense, and splice-region variants were predicted with bcftools. Missense variants identified in GS transcript reads were used to generate sample-specific protein sequences for each GS gene. Reference protein sequences were obtained from annotated genome protein FASTA files, and predicted missense substitutions were applied directly to the reference sequences at annotated positions using a custom Python script, resulting in one consensus protein sequence per sample per gene. Gene-specific protein FASTA files were aligned using CLC Sequence Viewer v8.0 (QIAGEN Digital Insights 2018).

## Results and Discussion

Failure to control *A. tuberculatus* populations with GA has been reported multiple times in recent years across Illinois farms, raising concerns about the potential evolution of resistance. Although escapes at the labeled recommended rate are often observed, resistance confirmation is challenging due to the significant effect of environmental conditions and weed size on the efficacy of GA, meaning that variable weed control can be expected if the labeled-recommended conditions of temperature, light, humidity, and weed size are not ideal at application. According to the International Survey of Herbicide-Resistant Weeds (Heap 2005), confirmation of resistance requires the fulfillment of four criteria: 1) the weed population must meet the formal definition of resistance by the Weed Science Society of America (WSSA); (2) there should be differential responses between the suspected-resistant and a known susceptible population of the same species across a wide range of herbicide rates; 3) the resistance heritability should be confirmed in the second generation; and 4) the weed species must be correctly identified. Based on these criteria, we conducted a series of field, greenhouse, and laboratory experiments to investigate four independent cases of GA control failure involving *A. tuberculatus* populations from Illinois fields.

### Greenhouse Whole-plant Dose-Response Assays

Initial greenhouse screenings carried out using progeny from field-collected seeds showed a range of 15 to 35% total survivorship among the suspected-resistant populations compared to 5% or lower survivorship in the herbicide-susceptible WUS population (data not shown). A preliminary dose-response study using progeny from field-collected seeds revealed that CAR exhibited an approximately two- to four-fold greater survival rate relative to WUS (data not shown). These preliminary observations prompted the development of a second generation of each population by crossing resistant plants, and subsequent dose-response experiments were conducted to confirm resistance is heritable.

In these assays, susceptible WUS plants were completely controlled 21 DAA by 371.1 g ai ha⁻¹ of GA (Table 1; Figure 2). The estimated GA rate required to reduce plant survival by 50% (LD₅₀) in the WUS population ranged from 66.3 to 170.2 g ai ha⁻¹ of GA across experimental runs, and the rate required to reduce dry biomass of WUS plants by 50% (GR₅₀) ranged from 41.5 to 115.9 g ai ha⁻¹ (Table 1). Based on plant survival, the estimated LD₅₀ values were 225.2, 360.1, and 167.3 g ai ha⁻¹ of GA for CAR, SDY, and FRA populations, respectively (Table 1; Figure 2). Estimated GR₅₀ values based on dry biomass were 115.1, 178.1, and 150.7 g ai ha⁻¹ of GA for CAR, SDY, and FRA populations, respectively. These represent a 3.4-, 2.1-, and 2.2-fold increase in LD₅₀ for CAR, SDY, and FRA, respectively, and a 2.8-, 1.9-, and 1.3-fold increase in GR₅₀, relative to the WUS population. The M01 population did not survive GA rates greater than 138.9 g ai ha⁻¹ across experimental runs, with only a few plants surviving at higher application rates; therefore, we were unable to model a dose-response curve of this population. Among the populations tested, M01 was the most sensitive to GA; however, preliminary screenings indicated survival at 370.1 ai ha⁻¹ of GA. Therefore, M01 was included in the RNA-Seq study, as discussed below.

**Table 1.** Parameter estimates from log-logistic regression models describing the responses of suspected-resistant and susceptible *Amaranthus tuberculatus* populations to glufosinate-ammonium in a greenhouse-based dose-response study 21 days after application.

| Population | Log-logistic regression model | Whole plant dose-response (g ai ha<sup>-1</sup>) |  |  |  |  |
| --- | --- | --- | --- | --- | --- | --- |
|  |  | Parameter estimates |  |  |  |  |
|  |  | b<sup>a</sup> (± SE) | c<sup>a</sup> (± SE) | d<sup>a</sup> (± SE) | e<sup>a</sup> (± SE) | LD<sub>50</sub>/GR<sub>50</sub><sup>b</sup> |
| ----- *Plant survival* ----- |  |  |  |  |  |  |
| CAR | 3-parameter | 1.59 (0.33) | - | 0.98 (0.09) | 225.22 (42.62) | 3.4 |
| WUS | log-logistic | 1.16 (0.45) | - | 1.00 (0.09) | 66.30 (35.53) |  |
| SDY | 3-parameter | 4.02 (0.94) | - | 1.00 (0.06) | 360.07 (27.81) | 2.1 |
| WUS | log-logistic | 2.03 (0.43) | - | 0.99 (0.08) | 170.18 (24.94) |  |
| FRA | 3-parameter | 4.21 (0.40) | - | 1.00 (0.02) | 167.32 (4.05) | 2.2 |
| WUS | log-logistic | 2.82 (1.01) | - | 1.00 (0.02) | 76.90 (19.00) |  |
| ----- *Dry biomass* ----- |  |  |  |  |  |  |
| CAR | 4-parameter | 2.76 (1.38) | 0.27 (0.07) | 2.23 (0.09) | 115.08 (15.53) | 2.8 |
| WUS | log-logistic | 1.7 (1.51) | 0.21 (0.07) | 2.77 (0.09) | 41.45 (44.37) |  |
| SDY | 4-parameter | 3.43 (0.90) | 0.27 (0.09) | 2.57 (0.15) | 178.09 (14.20) | 1.9 |
| WUS | log-logistic | 3.17 (1.92) | 0.28 (0.08) | 3.40 (0.15) | 95.73 (24.19) |  |
| FRA | 4-parameter | 8.36 (0.35) | 0.21 (0.01) | 2.21 (0.01) | 150.72 (0.81) | 1.3 |
| WUS | log-logistic | 9.55 (0.63) | 0.20 (0.01) | 2.02 (0.01) | 115.94 (13.76) |  |

<sup>a</sup> b is the slope around the effective rate, c is the lower asymptote, d is the upper asymptote, and e is the effective rate causing 50% reduction in response.
<sup>b</sup> LD<sub>50</sub> is the herbicide rate causing 50% mortality; GR<sub>50</sub> is the herbicide rate causing 50% biomass reduction relative to the susceptible population.

**Figure 2.**
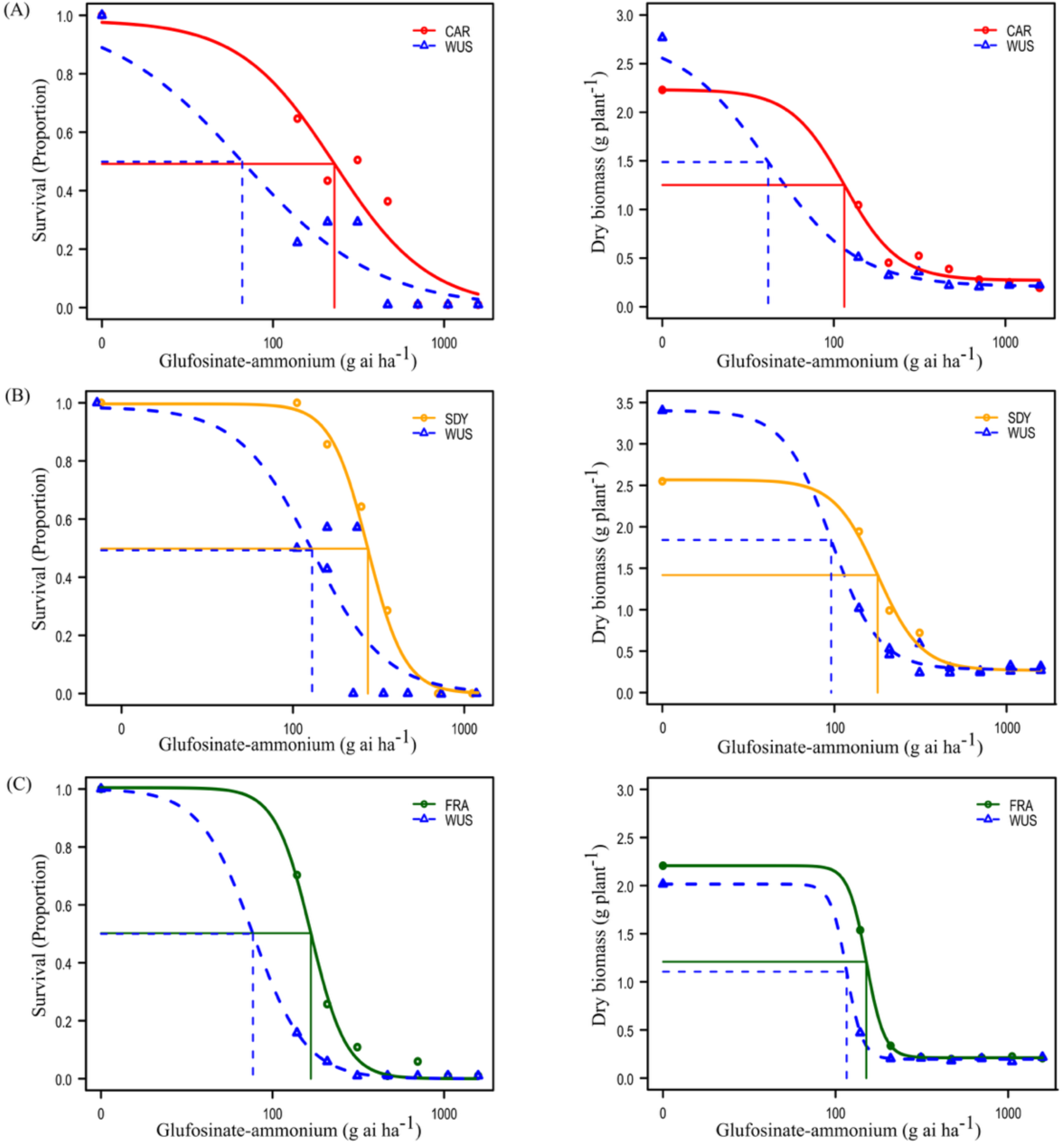
Glufosinate-ammonium dose-response of *Amaranthus tuberculatus* populations from (A) CAR, (B) SDY, and (C) FRA, based on plant survival (left panels) and dry biomass per plant (right panels), compared to the herbicide-susceptible population (WUS). Vertical lines indicate the estimated GR₅₀ values.

In the genus *Amaranthus*, GA resistance has only been documented previously in *A. palmeri*. In 2022, three *A. palmeri* populations from Arkansas were confirmed resistant to GA, with resistance ratios ranging from 5.1- to 27.4-fold relative to susceptible populations (Priess et al. 2022). In the same year, a GA-resistant *A. palmeri* population from Missouri was reported with LD₅₀ values indicating a 4.1-fold resistance in the field-collected population and a 6.1-fold resistance reported in the progeny (Noguera et al. 2022). More recently, an *A. palmeri* population from North Carolina exhibited LD₅₀ values ranging from 1.5- to 2.3-fold compared to a susceptible population (Jones et al. 2024). Although resistance ratios were slightly lower in our experiments, the observation of reduced sensitivity in progeny derived from field populations compared to susceptible populations confirms that GA resistance is a heritable trait. Given that GA resistance in *A. palmeri* is generally higher and known to be conferred by target-site mechanisms, it is possible that the lower level of resistance here could be explained by the early-stage evolution of complex nontarget-site (NTS) resistance.

### Field Dose-Response Study

The efficacy of GA in field applications is strongly influenced by prevailing environmental conditions, which are not easily replicated in greenhouse studies. To evaluate the response to GA under field conditions, we performed a field study at the location where the CAR population was originally collected. Results showed that GA provided less than 90% control 24 DAA when applied at one-half the labeled rate, and up to 20% of plants survived at the labeled rate (371.1 g ai ha⁻¹; Liberty® ULTRA Herbicide) (Figure 3). Complete control was only achieved with GA at twice the labeled rate, consistent with greenhouse dose-response results, with a single escape at four times the labeled rate (Figure 3). Comparable reductions in control (<90%) are typically found with applications on larger *Amaranthus* spp. plants (>15 cm). In this study, treatments were applied to plants up to 12.5 cm under optimal environmental conditions, including high relative humidity (71%) and temperature (31 °C) that are known to enhance GA efficacy. In addition, no shading from crop plants was present at the time of application, allowing for thorough coverage of *A. tuberculatus* plants. Although survival levels may appear low in a research context, they are sufficient to promote the spread of resistance in the subsequent growing seasons, particularly under continued GA selection pressure.

**Figure 3.**
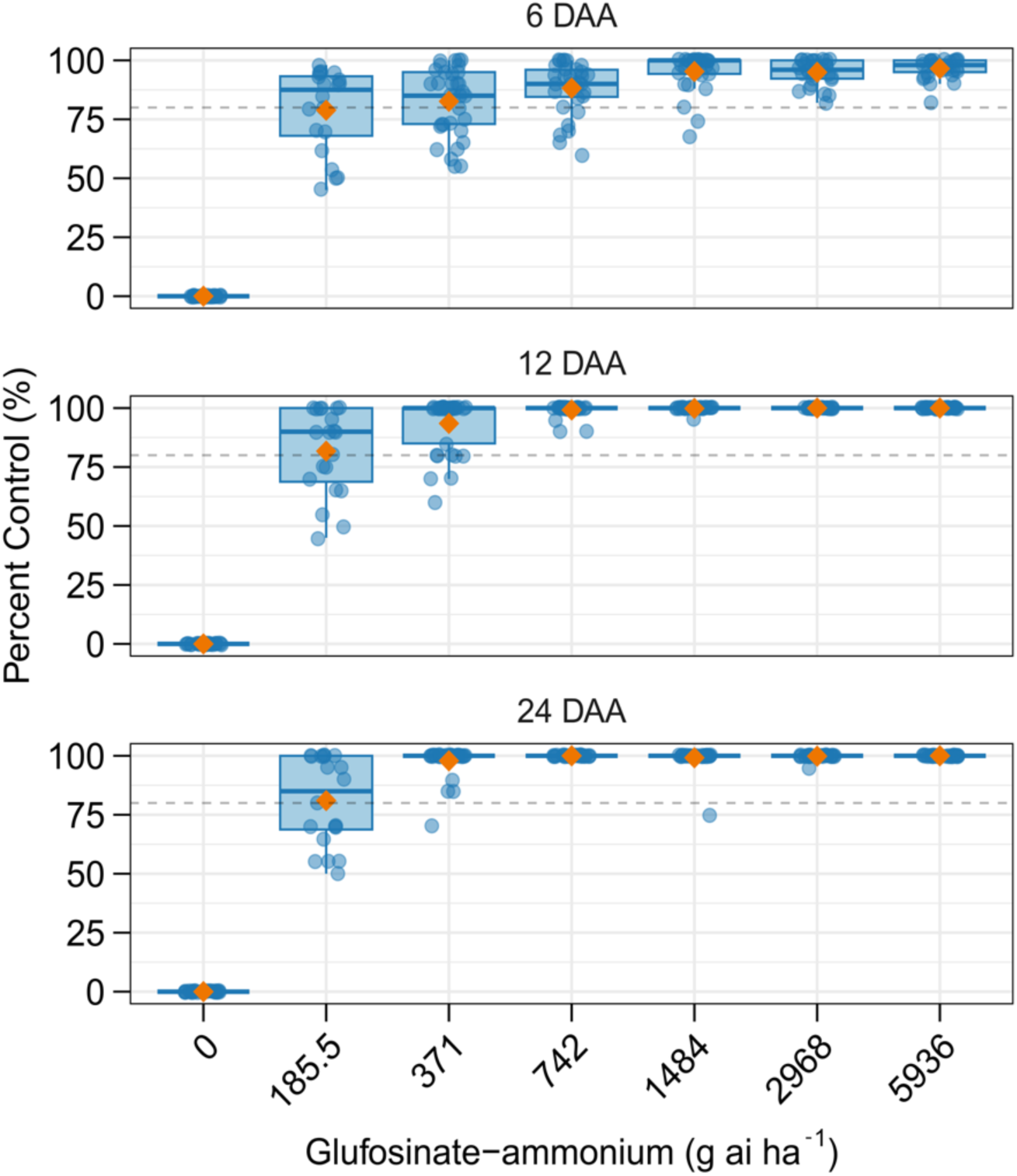
Boxplots showing percent control of the suspected-resistant *Amaranthus tuberculatus* population from Carroll County, Illinois, in response to increasing rates of glufosinate-ammonium in a field dose-response study at 6, 12, and 24 days after application. The vertical dashed line indicates the 80% control threshold. Blue points represent individual plants, and orange triangles denote the mean percentage of control at each herbicide rate.

### Multiple Herbicide Site-of-Action Screening

To identify alternative management options for the suspected GA-resistant populations, each population was screened with multiple herbicides representing different SOA groups. Glyphosate reduced biomass by ≥70% in the CAR and SDY populations at the 1× rate or higher; however, biomass reduction in the M01 population was <50% at the 1× rate relative to the nontreated control (Supplementary Figure S1). The susceptible control, WUS, exhibited >80% biomass reduction at the 1× rate. Compared to BRC, the FRA population exhibited reduced sensitivity to glyphosate, with plants surviving all three application rates and complete control not achieved at any rate (Supplementary Figure S2). In *A. tuberculatus*, resistance to glyphosate was first reported in 2005 and is now present in 20 U.S. states (Heap 2025). Typically, in *Amaranthus* species, glyphosate resistance arises through amplification of the target gene encoding enolpyruvyl shikimate-3-phosphate synthase (*EPSPS*), and in Illinois, approximately 91% of glyphosate-resistant *A. tuberculatus* populations exhibit *EPSPS* gene amplification (Chatham et al. 2015). Reduced glyphosate translocation and the Pro106Ser substitution in the *EPSPS* gene have also been reported in several *A. tuberculatus* populations, with some populations harboring both TS and NTS resistance mechanisms (Murphy et al. 2019; Nandula et al. 2013).

For atrazine, CAR and SDY were the least affected, exhibiting relatively high biomass (>65%) compared with nontreated plants, with CAR surviving the 3× rate (Supplementary Figure S1). In contrast, biomass reduction for M01 and WUS exceeded 80% at the 1× rate. For the FRA population, complete control with atrazine was achieved only at the 3× rate (Supplementary Figure S2). The photosystem II (PS II)-inhibiting herbicides was the first SOA group for which *A. tuberculatus* evolved resistance (Anderson et al. 1996). Resistance to PS II inhibitors can arise through a TS mutation in the D1 protein, conferring high levels of resistance, or through glutathione *S*-transferase (GST)-mediated enhanced metabolic detoxification, which typically confers more moderate resistance (Evans Jr et al. 2017).

CAR, SDY, and WUS exhibited >80% biomass reduction with fomesafen at the 1× rate, whereas M01 showed <65% biomass reduction at the same rate (Supplementary Figure S1). For the FRA population, complete control with the PPO-inhibiting herbicides fomesafen and lactofen was not achieved at any rate, although biomass was significantly reduced at the 3× rate (Supplementary Figure S2). Similarly, imazethapyr failed to control either FRA or BRC at any rate, consistent with the historically widespread resistance to acetolactate synthase (ALS)-inhibiting herbicides in Illinois *A. tuberculatus* populations, given that BRC was originally collected in 2002.

CAR, SDY, and WUS biomass reduction ≥80% was observed at the 1× and 3× rates of 2,4-D, although lower reduced biomass reduction was observed for SDY (70%) and M01 (50%) at the 1× rate. For FRA and BRC, complete control was achieved at all application rates of 2,4-D and paraquat (0.5×, 1×, and 3×). Overall, these results indicate that several populations exhibit reduced sensitivity to herbicides across multiple SOA groups, with responses varying by population. Management strategies should therefore be tailored to the populations present within individual fields. Based on these screenings, 2,4-D remained an effective herbicide option in this study, providing consistent control across most populations evaluated.

### Transcriptomes of A. tuberculatus populations

We next examined global transcriptome profiles to explore gene expression patterns potentially associated with resistance. Principal component analysis (PCA) of global transcriptome data showed clear separation of populations along PC1, which explained 21.6% of the total variance (Figure 4A). The susceptible populations (WUS and BRC) clustered together and were separated from the suspected resistant populations (CAR, SDY, FRA, and M01), indicating broad differences in overall gene expression profiles associated with resistance status. PC2 explained an additional 9.0% of the variance and captured further transcriptomic heterogeneity among populations, particularly within the suspected resistant group. Venn diagram analyses revealed that relatively few DEGs were shared among the suspected resistant populations (Figure 4B). Most DEGs were unique to individual populations, with only a small subset overlapping across two or more populations, indicating limited convergence at the individual gene level among the suspected resistant populations. Consistent with the PCA results, clustering of DEGs in the heat map showed that the susceptible populations WUS and BRC formed a distinct cluster despite originating from different states, suggesting that their similarity in expression patterns is unlikely to reflect geographic proximity (Figures 1 and 4C). Clustering was also observed among the suspected resistant populations CAR, SDY, and FRA, with SDY showing an intermediate expression profile that partially overlapped with the susceptible populations. Together, these results indicate that although suspected resistant populations show similarity in global gene expression patterns, this similarity is more likely driven by broad, coordinated shifts in transcriptomic profiles rather than by shared differentially expressed genes. This pattern suggests that suspected resistant populations may share common physiological or regulatory responses associated with resistance, while the specific gene-level changes contributing to these responses remain largely population-specific.

**Figure 4.**
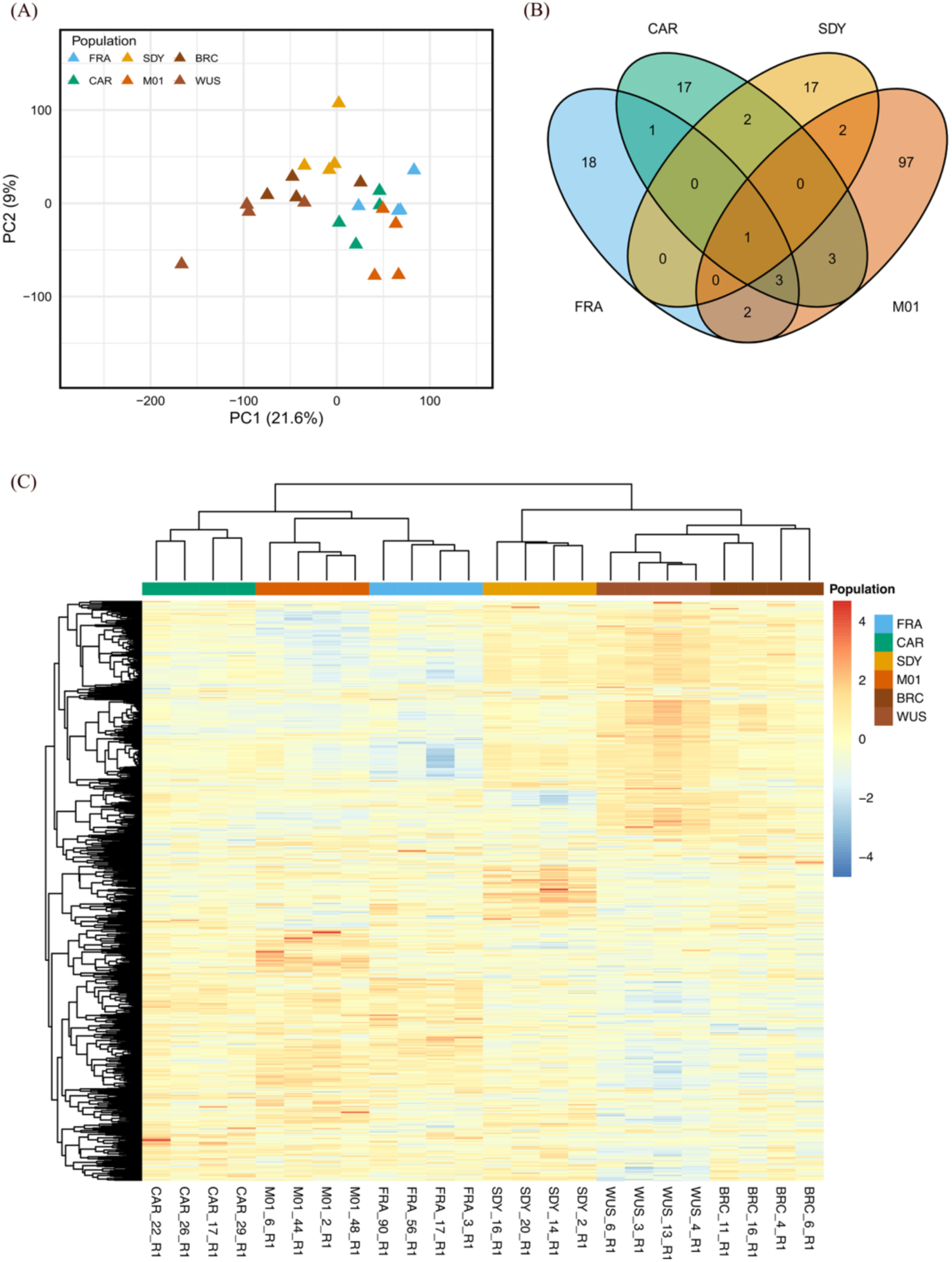
Summary of gene expression profiles of suspected-resistant *Amaranthus tuberculatus* populations (CAR, FRA, SDY, M01) and susceptible populations (WUS, BRC). (A) Principal Component Analysis based on global transcriptome profiles. (B) Venn diagrams of shared differentially expressed genes (DEGs) between resistant populations compared individually with both susceptible populations. (C) Heat map of 456 DEGs identified across comparisons with WUS and BRC.

To explore candidate genes associated with GA resistance, we focused on DEGs consistently identified in comparisons with both WUS and BRC. The analysis revealed 268, 256, 503, and 618 DEGs when CAR, SDY, FRA, and M01 were compared with WUS, and 39, 44, 33, and 155 DEGs when compared with BRC, respectively (Supplementary Tables S3–S10). Restricting the analysis to DEGs shared in comparisons with both susceptible populations yielded 27 (CAR), 22 (SDY), 25 (FRA), and 108 (M01) genes (Figure 4B; Supplementary Table S11). One DEG (*AmaTu_RefChr07g124540*), annotated as a TolB-like protein, was consistently upregulated and shared across all four suspected resistant populations. Additional BLAST searches suggested similarity to WD40-like β-propeller proteins, which typically serve as molecular hubs by signaling and enhancing interactions between multiple proteins, from forming complexes with transcription factors to regulating stress-responsive genes to being involved in histone modifications and chromatin remodeling (Meng et al. 2024; Shoeva et al. 2023; Suganuma et al. 2008; Yuan et al. 2019). Additional DEGs shared among CAR, FRA, and M01 included an F-box phloem protein 2 (PP2; *AmaTu_RefChr06g107540*) that was upregulated in all three populations and a 30S ribosomal protein (*AmaTu_RefChr08g13286*0) that was downregulated in all three.

Beyond these shared DEGs, each population had distinct sets of putative resistance-associated genes. In CAR, these included upregulated genes such as UDP-glycosyltransferases, glutaredoxin C9, 1-aminocyclopropane-1-carboxylate synthase/oxidase, and a WRKY transcription factor, as well as downregulated genes such as 4-hydroxy-tetrahydrodipicolinate synthase, 1-acyl-sn-glycerol-3-phosphate acyltransferase 5, and a MYC transcription factor. In addition to those, genes involved in stress signaling, as well as genes encoding structural proteins, were also identified (Supplementary Tables S3, S4, and S11).

In SDY, genes identified included upregulated genes such as glutathione *S*-transferases, 12-oxophytodienoate reductase 3-like, 3-oxo-Delta(4,5)-steroid 5-beta-reductase-like, and a GLK1-like transcription factor, and downregulated genes including a cytochrome P450 (*CYP82D47*), and a MYC-like transcription factor (Supplementary Tables S5, S6, and S11). In FRA, interesting genes included an upregulated cytochrome P450 (*CYP81E8*) gene and several upregulated transcription factors (A-3-like, WRKY, and NF-X1-type zinc finger proteins) (Supplementary Tables S7, S8, and S11). Two upregulated cytochrome P450 genes were identified in M01, one also annotated as *CYP81E8* but located on a different chromosome than the one identified in FRA, and a *CYP71A26*. An ABC transporter gene was also upregulated, as well as several other transporter and transcription factor genes were differentially expressed in this population (Supplementary Tables S9–S11).

Cytochrome P450s, GSTs, and glycosyltransferases are frequently implicated in multigenic, metabolism-based herbicide resistance in weeds, and the presence of these gene families among the DEGs is consistent with such mechanisms in the *A. tuberculatus* populations in this study (Bobadilla and Tranel 2024a; Concepcion et al. 2021; Dimaano and Iwakami 2021; Evans Jr et al. 2017). The distinct set of DEGs among populations would suggest that different resistance mechanisms may underlie reduced GA sensitivity. Nevertheless, it should also be noted here that most of these populations appear to be resistant to other herbicide chemistries, as seen earlier with the SOA screening results. This could indicate that some of the identified resistance genes may not be directly linked to GA resistance but rather associated with resistance to other herbicides. For instance, the cytochrome P450 gene *CYP81E8* identified in M01 has previously been implicated in resistance to 2,4-D in two *A. tuberculatus*, with phylogenetic evidence suggesting a shared evolutionary origin of the resistant alleles in these populations (Giacomini et al. 2020). Therefore, it is possible that some of the identified genes contribute to cross-resistance to GA. This scenario is not uncommon with multiple-resistant weed populations and could also explain why the populations in this study are now gradually evolving resistance to GA (Bobadilla and Tranel 2024b).

To gain additional insight into the molecular basis of resistance, we performed GO enrichment analysis of the DEGs for each population (Supplementary Tables S12–S15; Figure 5). In the CAR population, DEGs were enriched for stress response, defense regulation and salicylic acid and abscisic acid metabolism. Differentially expressed gene products were predicted to be located in the cytoplasm, nucleus, ribosomes, and plastids with functions spanning DNA/RNA binding, transcription regulation, and metabolic modifications. In SDY, genes were significantly associated with regulation of gene expression, including transcription factor activity, RNA binding, and mRNA stability. Genes also showed enrichment for secondary metabolism and stress-response pathways. The gene products were predicted to be located predominantly in the cytosol, ribosomal units, chloroplasts and plastids, and peroxisomes, which could indicate functions in photosynthetic processes and oxidative related metabolism.

**Figure 5.**
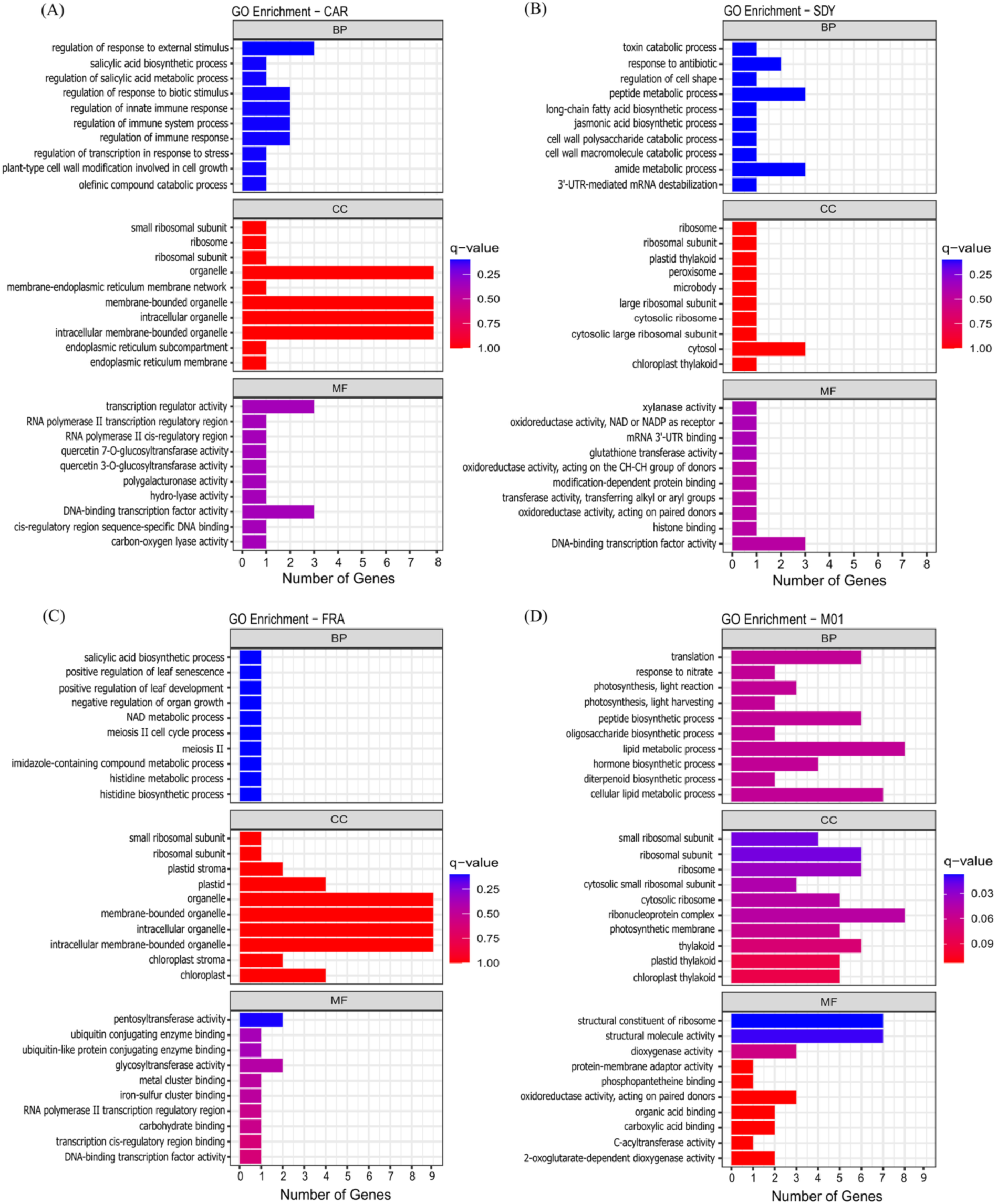
Gene Ontology (GO) enrichment analysis of suspected-resistant *Amaranthus tuberculatus* populations, showing biological process (BP), cellular component (CC), and molecular function (MF) terms for CAR (A), SDY (B), FRA (C), and M01 (D).

In FRA, DEGs were enriched for nitrogen and secondary metabolism, stress and defense signaling, and growth regulation, and were predicted to be located in the chloroplasts, plastids, and ribosomes. Functionally, these genes are likely involved in transcription regulation, protein modifications, and other enzymatic activities. In M01, enriched terms pointed to hormone, lipid, peptide, and secondary metabolism of other metabolites. Other enriched terms also indicate activity in translation and ribosomal processes, with most gene products being located in ribosomes and in the chloroplast. Overall, the molecular function of these genes appears to be related to molecule binding, catalytic processes, and molecule transport activities.

### Identification of Glutamine Synthetase Homologs

A variable number of GS genes can be found in plant species. In diploid species, the cytosolic GS (*GS1*) is typically encoded by multiple genes, while the plastidic GS (*GS2*) is in general encoded by a single nuclear gene (Cánovas et al. 2007; Valderrama-Martín et al. 2022). However, some diploid species such as barrel medic (*Medicago truncatula*) harbor a second *GS2* copy (Seabra et al. 2010), and more recent studies also reported the presence of at least two *GS2* copies in *A. palmeri*, with evidence suggesting that they can contribute differentially to GA resistance (Carvalho-Moore et al. 2025; Noguera et al. 2024, 2022). Similarly, in our study, two copies of *GS2* were identified in *A. tuberculatus* (Supplementary Figure S3), and transcriptome data confirmed that both copies are expressed. Preliminary transcriptomic evidence suggests that *GS2.1* and *GS2.2* might not be equally expressed; however, this should be further confirmed using real-time quantitative PCR in a future study. To explore the number of *GS2* copies in other *Amaranthus* species, we obtained *GS1* and *GS2* sequences of multiple species from publicly available genomic resources. Interestingly, the two plastidic GS copies (*GS2.1* and *GS2.2*) were detected in all species except for *A. hypochondriacus* (Supplementary Figure S3). Given that the *A. hypochondriacus* assembly remains at the scaffold level, it is possible that additional GS copies exist in its genome but are unresolved in the current genome version. In all *Amaranthus* species having two *GS2* copies, *GS2.1* and *GS2.2* were positioned on the same chromosome in close physical proximity. These observations suggest that the presence of multiple *GS2* copies may be far more common within the *Amaranthus* genus than previously anticipated.

In previous reports of GA resistance in *A. palmeri*, overexpression of *GS2* was proposed as the primary resistance mechanism (Carvalho-Moore et al. 2022; Noguera et al. 2022). To assess whether this mechanism contributed to resistance in the populations studied herein, we explored the expression of both *GS1* and *GS2* copies in resistant and susceptible populations. None of the four populations showed significant differences in expression of any of the *GS1* or *GS2* copies relative to the susceptible WUS or BRC populations. We further examined the coding sequences of both the *GS1* and *GS2* copies to identify mutations that could confer GA resistance. A summary of all identified missense variants across the four genes is provided in Supplementary Table S16. In total, four missense SNPs were identified in *GS2.1*, thirteen in *GS2.2*, three in *GS1.1*, and seven in *GS1.2* (Supplementary Figures S4-S7). None of these SNPs occurred within predicted catalytic domains of *GS*, as recently summarized by Porri et al. 2025, and therefore they are unlikely to be involved in GA resistance.

Together, our findings provide evidence for the evolution of GA resistance in populations of *A. tuberculatus* in Illinois. While currently we only have sufficient evidence to confirm resistance in the CAR population, reduced sensitivity and the upregulation of multiple metabolism-related genes in SDY, FRA, and M01 suggests that GA resistance is evolving in these populations and may increase in prevalence under continued selection pressure. The observed expression of metabolism-related genes also signals multigenic mechanisms with the potential for cross-resistance across herbicide chemistries. Given that all four populations showed high survival rates to multiple herbicide SOA groups, it is likely that some of the identified genes contribute to broader resistance and possibly cross-resistance to GA. This is of particular concern given that *A. tuberculatus* populations in Illinois are already known to have evolved resistance to herbicides spanning seven SOA groups. Importantly, our results demonstrated that 2,4-D remains a reliable control option for most populations, while atrazine, glyphosate, lactofen, fomesafen, and imazethapyr provided less *A. tuberculatus* control. From a management perspective, these findings highlight the need for more proactive, diversified weed control strategies in Midwestern fields. Integrating tactics such as narrow row cropping systems, cover crops, and harvest weed seed control is essential for preserving herbicide usefulness and long-term management of *A. tuberculatus* as resistance continues to evolve.

## Author contributions

L.M. and C.B.R. collected seeds from field populations. I.W.N., C.B.R., L.M., and E.L. conducted greenhouse experiments. I.W.N., L.M., R.S., and A.J.L. carried out the field study. I.W.N. performed the statistical and bioinformatics analyses. I.W.N., C.B.R., E.L., and L.M drafted the original manuscript. All authors reviewed and edited the manuscript. P.J.T., A.G.H., and K.G. acquired funding, conceptualized the study, and supervised the research.

## Acknowledgments

This work was supported by the Illinois Soybean Checkoff Program, the Soy Checkoff, and Hatch accession no. 7000961 and by United Soybean Board prime agreement no. 25-210-S-B-2-A, sub-agreement no. 40005185-202.

## References

Alexa A, Rahnenführer J (2009) Gene set enrichment analysis with topGO. Bioconductor Improv 27:776

Anderson DD, Roeth FW, Martin AR (1996) Occurrence and control of triazine-resistant common waterhemp (*Amaranthus rudis*) in field corn (*Zea mays*). Weed Technol 10:570–575, 10.1017/S0890037X00040458

Anonymous (2024) Liberty® ULTRA herbicide product label. EPA Reg. No. 7969-500. https://www.cdms.telusagcg.com/ldat/ldJGM003.pdf. Accessed on December 6, 2025.

Babraham Bioinformatics. 2019. FastQC: A quality control tool for high throughput sequence data. Version 0.11.9. https://www.bioinformatics.babraham.ac.uk/projects/fastqc/. Accessed: January 14, 2026

Baker NT, Stone WW (2015). Estimated annual agricultural pesticide use for counties of the conterminous United States, 2008-12 (No. 907). US Geological Survey.

Bobadilla LK, Tranel PJ (2024a) Identification of candidate genes involved with dicamba resistance in waterhemp (*Amaranthus tuberculatus*) via transcriptomics analyses. Weed Sci 72:125–136, 10.1017/wsc.2023.73

Bobadilla LK, Tranel PJ (2024b) Predicting the unpredictable: the regulatory nature and promiscuity of herbicide cross resistance. Pest Manag Sci 80:235–244, 10.1002/ps.7728

Brown AK, Farenhorst A (2024) Quantitation of glyphosate, glufosinate, and AMPA in drinking water and surface waters using direct injection and charged-surface ultra-high performance liquid chromatography-tandem mass spectrometry. Chemosphere 349:140924, 10.1016/j.chemosphere.2023.140924

Cánovas FM, Ávila C, Cantón FR, Cañas RA, de la Torre F (2007) Ammonium assimilation and amino acid metabolism in conifers. J Exp Bot 58:2307–2318. 10.1093/jxb/erm051

Cantalapiedra CP, Hernández-Plaza A, Letunic I, Bork P, Huerta-Cepas J (2021) eggNOG-mapper v2: functional annotation, orthology assignments, and domain prediction at the metagenomic scale. Mol Biol Evo 38:5825–5829, 10.1093/molbev/msab293

Carvalho-Moore P, Borgato EA, Cutti L, Porri A, Meiners I, Lerchl J, Norsworthy JK, Patterson EL (2025) A rearranged *Amaranthus palmeri* extrachromosomal circular DNA confers resistance to glyphosate and glufosinate. Plant Cell 37: koaf069, 10.1093/plcell/koaf069

Carvalho-Moore P, Norsworthy JK, González-Torralva F, Hwang J-I, Patel JD, Barber LT, Butts TR, McElroy JS (2022) Unraveling the mechanism of resistance in a glufosinate-resistant Palmer amaranth (*Amaranthus palmeri*) accession. Weed Sci 70:370–379, 10.1017/wsc.2022.31

Chatham LA, Wu C, Riggins CW, Hager AG, Young BG, Roskamp GK, Tranel PJ (2015) *EPSPS* gene amplification is present in the majority of glyphosate-resistant Illinois waterhemp (*Amaranthus tuberculatus*) populations. Weed Technol 29:48–55, 10.1614/WT-D-14-00064.1

Coetzer E, Al-Khatib K, Loughin TM (2001) Glufosinate efficacy, absorption, and translocation in amaranth as affected by relative humidity and temperature. Weed Sci 49:8–13, 10.1614/0043-1745(2001)049[0008:GEAATI]2.0.CO;2

Concepcion JCT, Kaundun SS, Morris JA, Hutchings SJ, Strom SA, Lygin AV, Riechers DE (2021) Resistance to a nonselective 4-hydroxyphenylpyruvate dioxygenase-inhibiting herbicide via novel reduction–dehydration–glutathione conjugation in *Amaranthus tuberculatus*. New Phytol 232:2089–2105, 10.1111/nph.17708

Danecek P, McCarthy A (2017) BCFtools/csq: haplotype-aware variant consequences. Bioinformatics 33:2037–2039

Dimaano NG, Iwakami S (2021) Cytochrome P450-mediated herbicide metabolism in plants: current understanding and prospects. Pest Manag Sci 77:22–32, 10.1002/ps.6040

Dobin A, Davis CA, Schlesinger F, Drenkow J, Zaleski C, Jha S, Batut P, Chaisson M, Gingeras TR (2013) STAR: ultrafast universal RNA-seq aligner. Bioinform 29:15–21, 10.1093/bioinformatics/bts635

Evans Jr AF, O’Brien SR, Ma R, Hager AG, Riggins CW, Lambert KN, Riechers DE (2017) Biochemical characterization of metabolism-based atrazine resistance in *Amaranthus tuberculatus* and identification of an expressed GST associated with resistance. Plant Biotechnol J 15:1238–1249, 10.1111/pbi.12711

Ewels P, Magnusson M, Lundin S (2016) MultiQC MK Summarize analysis results for multiple tools and samples in a single report. Bioinform 32: 3047–3048, 10.1093/bioinformatics/btw354

Giacomini DA, Patterson EL, Küpper A, Beffa R, Gaines TA, Tranel PJ (2020) Coexpression clusters and allele-specific expression in metabolism-based herbicide resistance. Genome Biol and Evol 12:2267–2278, 10.1093/gbe/evaa191

Haarmann JA, Young BG, Johnson WG (2020) Control of waterhemp (*Amaranthus tuberculatus*) regrowth after failed applications of glufosinate or fomesafen. Weed Technol 34:794–800, 10.1017/wet.2020.58

Heap I (2005) Criteria for confirmation of herbicide-resistant weeds. Weed Science 53:652–658

Heap I (2025) International Herbicide-Resistant Weed Database. https://www.weedscience.org/. Accessed July 21, 2025

Jones EA, Dunne JC, Cahoon CW, Jennings KM, Leon RG, Everman WJ (2024) Confirmation and inheritance of glufosinate resistance in an *Amaranthus palmeri* population from North Carolina. Plant-Environ Interact 5:e10154, 10.1002/pei3.10154

Kolde R, Kolde MR (2015) Package ‘pheatmap’. R package version 1.0.12. https://cran.r-project.org/web/packages/pheatmap/. Accessed September 9, 2025

Kumar S, Stecher G, Suleski M, Sanderford M, Sharma S, Tamura K (2024) MEGA12: Molecular Evolutionary Genetic Analysis version 12 for adaptive and green computing. Mol Biol Evol 41:msae263. 10.1093/molbev/msae263

Kumaratilake AR, Preston C (2005) Low temperature reduces glufosinate activity and translocation in wild radish (*Raphanus raphanistrum*). Weed Sci 53:10–16, 10.1614/WS-03-140R

Landau C, Bradley K, Burns E, DeWerff R, Dobbels A, Essman A, Flessner M, Gage K, Hager A, Jhala A (2025) Weather and glufosinate efficacy: a retrospective analysis looking forward to the changing climate. Weed Sci 73:e32, 10.1017/wsc.2024.101

Letunic I, Bork P (2024) Interactive Tree of Life (iTOL) v6: recent updates to the phylogenetic tree display and annotation tool. Nucleic Acids Res 52:W78–W82. 10.1093/nar/gkae268

Liao Y, Smyth GK, Shi W (2014) featureCounts: an efficient general purpose program for assigning sequence reads to genomic features. Bioinform 30:923–930, 10.1093/bioinformatics/btt656

Lightfoot DJ, Jarvis DE, Ramaraj T, Lee R, Jellen EN, Maughan PJ (2017) Single-molecule sequencing and Hi-C-based proximity-guided assembly of amaranth (*Amaranthus hypochondriacus*) chromosomes provide insights into genome evolution. BMC Biol, 15(1):74, 10.1186/s12915-017-0412-4

Lingenfelter D (2024) Follow these steps to avoid glufosinate resistance. https://www.farmprogress.com/commentary/follow-these-steps-to-avoid-glufosinate-resistance. Accessed: September 30, 2025

Meng L, Su H, Qu Z, Lu P, Tao J, Li H, Zhang J, Zhang W, Liu N, Cao P, Jin J (2024) Genome-wide identification and analysis of WD40 proteins reveal that NtTTG1 enhances drought tolerance in tobacco (*Nicotiana tabacum*). BMC Genomics 25:133, 10.1186/s12864-024-10022-w

Meyer CJ, Norsworthy JK (2020) Timing and application rate for sequential applications of glufosinate are critical for maximizing control of annual weeds in LibertyLink® Soybean. Int J Agron 2020:9145370, 10.1155/2020/9145370

Murphy BP, Larran AS, Ackley B, Loux MM, Tranel PJ (2019) Survey of glyphosate-, atrazine-and lactofen-resistance mechanisms in Ohio waterhemp (*Amaranthus tuberculatus*) populations. Weed Sci 67:296–302, 10.1017/wsc.2018.91

Nandula VK, Ray JD, Ribeiro DN, Pan Z, Reddy KN (2013) Glyphosate resistance in tall waterhemp (*Amaranthus tuberculatus*) from Mississippi is due to both altered target-site and nontarget-site mechanisms. Weed Sci 61:374–383, 10.1614/WS-D-12-00155.1

Noguera MM, Albert PS, Saski CA, Birchler JA, Porri A, Meiners I, Lerchl J, Roma-Burgos N (2024) Extrachromosomal DNA-mediated glutamine synthetase 2 (*GS2*) amplification enables glufosinate resistance in *Amaranthus palmeri*. bioRxiv:2024.12.13.628355, 10.1101/2024.12.13.628355.

Noguera MM, Porri A, Werle IS, Heiser J, Brändle F, Lerchl J, Murphy B, Betz M, Gatzmann F, Penkert M, Tuerk C, Meyer L, Roma-Burgos N (2022) Involvement of glutamine synthetase 2 (*GS2*) amplification and overexpression in *Amaranthus palmeri* resistance to glufosinate. Planta 256:57, 10.1007/s00425-022-03968-2

Okada E, Coggan T, Anumol T, Clarke B, Allinson G (2019) A simple and rapid direct injection method for the determination of glyphosate and AMPA in environmental water samples. Anal and Bioanal Chem 411:715–724, 10.1007/s00216-018-1490-z

Porri A, Sudhakar S, Noguera MM, Betz M, Lerchl J, Dayan FE, Norsworthy JK (2025) Analysis of glutamine synthetase target-site mutations and their role in endowing glufosinate-ammonium resistance. bioRxiv:2025.11.01.685562, 10.1101/2025.11.01.685562

Preston C (2024) Strategies for optimising glufosinate and tackling efficacy challenges. Canberra, Australia. Grains Research & Development Corporation. https://grdc.com.au/resources-and-publications/grdc-update-papers/tab-content/grdc-update-papers/2024/02/strategies-for-optimising-glufosinate-and-tackling-efficacy-challenges Accessed: January 3, 2026

Priess GL, Norsworthy JK, Godara N, Mauromoustakos A, Butts TR, Roberts TL, Barber T (2022) Confirmation of glufosinate-resistant *Palmer amaranth* and response to other herbicides. Weed Technol 36:368–372, 10.1017/wet.2022.21

QIAGEN Digital Insights (2018) CLC Sequence Viewer User Manual, version 8.0. QIAGEN, Aarhus, Denmark. https://resources.qiagenbioinformatics.com/manuals/clcsequenceviewer/current/User_Manual.pdf Accessed: January 12, 2026

Raiyemo DA, Cutti L, Patterson EL, Llaca V, Fengler K, Montgomery JS, Morran S, Gaines TA, Tranel PJ (2024) A phased chromosome-level genome assembly provides insights into the evolution of sex chromosomes in *Amaranthus tuberculatus*. bioRxiv:2024.05. 30.596720, 10.1101/2024.05.30.596720

Raiyemo DA, Montgomery JS, Cutti L, Abdollahi F, Llaca V, Fengler K, Lopez AJ, Morran S, Saski CA, Nelson DR, Patterson EL, Gaines TA, Tranel PJ (2025) Chromosome-level assemblies of *Amaranthus palmeri*, *Amaranthus retroflexus*, and *Amaranthus hybridus* allow for genomic comparisons and identification of a sex-determining region. Plant J, 121: e70027, 10.1111/tpj.70027

Ramsey R, Stephenson G, Hall J (2006) Effect of humectants on the uptake and efficacy of glufosinate in wild oat (*Avena fatua*) plants and isolated cuticles under dry conditions. Weed Sci 54:205–211, 10.1614/WS-05-051R.1

Risso D, Ngai J, Speed TP, Dudoit S (2014) Normalization of RNA-seq data using factor analysis of control genes or samples. Nat Biotechnol 32:896–902

Ritz C, Baty F, Streibig JC, Gerhard D (2015) Dose-Response analysis using R. PLoS One 10:e0146021, 10.1371/journal.pone.0146021

Robinson MD, McCarthy DJ, Smyth GK (2010) edgeR: a Bioconductor package for differential expression analysis of digital gene expression data. Bioinform 26:139–140, 10.1093/bioinformatics/btp616

Seabra AR, Vieira CP, Cullimore JV, Carvalho HG (2010) *Medicago truncatula* contains a second gene encoding a plastid-located glutamine synthetase exclusively expressed in developing seeds. BMC Plant Biol 10:183. 10.1186/1471-2229-10-183

Shoeva OY, Mukhanova MA, Zakhrabekova S, Hansson M (2023) Ant13 encodes regulatory factor WD40 controlling anthocyanin and proanthocyanidin synthesis in barley (Hordeum vulgare L.). J Agric Food Chem 71:6967–6977, 10.1021/acs.jafc.2c09051.

Simms D, Cizdziel PE, Chomczynski P (1993) TRIzol: A new reagent for optimal single-step isolation of RNA. Focus 15:532–535

Singh N, Sarangi D, Stahl L, Ikley J, Peters T (2024) Best practices for using glufosinate (Liberty) herbicide. https://blog-crop-news.extension.umn.edu/2024/05/best-practices-for-using-glufosinate.html#:~:text=Temperature:%20The%20Liberty%20label%20states,versus%20earlier%20in%20the%20season. Accessed: September 30, 2025

Steckel GJ, Wax LM, Simmons FW, Phillips WH (1997) Glufosinate efficacy on annual weeds is influenced by rate and growth stage. Weed Technol 11:484–488, 10.1017/S0890037X00045292

Suganuma T, Pattenden SG, Workman JL (2008) Diverse functions of WD40 repeat proteins in histone recognition. Genes Dev 22:1265–1268, 10.1101/gad.1676208

Takano HK, Beffa R, Preston C, Westra P, Dayan FE (2019) Reactive oxygen species trigger the fast action of glufosinate. Planta 249:1837–1849, 10.1007/s00425-019-03124-3

Takano HK, Beffa R, Preston C, Westra P, Dayan FE (2020a) A novel insight into the mode of action of glufosinate: how reactive oxygen species are formed. Photosynth Res 144:361–372, 10.1007/s11120-020-00749-4

Takano HK, Beffa R, Preston C, Westra P, Dayan FE (2020b) Physiological factors affecting uptake and translocation of glufosinate. J Agric Food Chem 68:3026–3032, 10.1021/acs.jafc.9b07046

Takano HK, Dayan FE (2020) Glufosinate-ammonium: a review of the current state of knowledge. Pest Manag Sci 76:3911–3925, 10.1002/ps.5965

Takano HK, Dayan FE (2021) Biochemical basis for the time-of-day effect on glufosinate efficacy against *Amaranthus palmeri*. Plants 10:2021, 10.3390/plants10102021

[USGS] U.S. Geological Survey (2018) Pesticide National Synthesis Project. https://water.usgs.gov/nawqa/pnsp/usage/maps/. Accessed: September 9, 2025

Valderrama-Martín JM, Ortigosa F, Ávila C, Cánovas FM, Hirel B, Cantón FR, Cañas RA (2022) A revised view on the evolution of glutamine synthetase isoenzymes in plants. A revised view on the evolution of glutamine synthetase isoenzymes in plants. Plant Journal 110(4):946–960, 10.1111/tpj.15712

Wang H, Xu D, Wang S, Wang A, Lei L, Jiang F, Yang B, Yuan L, Chen R, Zhang Y, Fan W (2023) Chromosome-scale *Amaranthus tricolor* genome provides insights into the evolution of the genus *Amaranthus* and the mechanism of betalain biosynthesis. DNA Res 30(1):dsac050, 10.1093/dnares/dsac050

Yuan F, Leng B, Zhang H, Wang X, Han G, Wang B (2019) A WD40-repeat protein from the recretohalophyte *Limonium bicolor* enhances trichome formation and salt tolerance in *Arabidopsis*. Front Plant Sci 10:1456, 10.3389/fpls.2019.01456

