## Supplemental Data 1 for "Confirmation and Transcriptomic Characterization of Glufosinate-ammonium Resistance in Waterhemp (*Amaranthus tuberculatus*) Populations from Illinois"

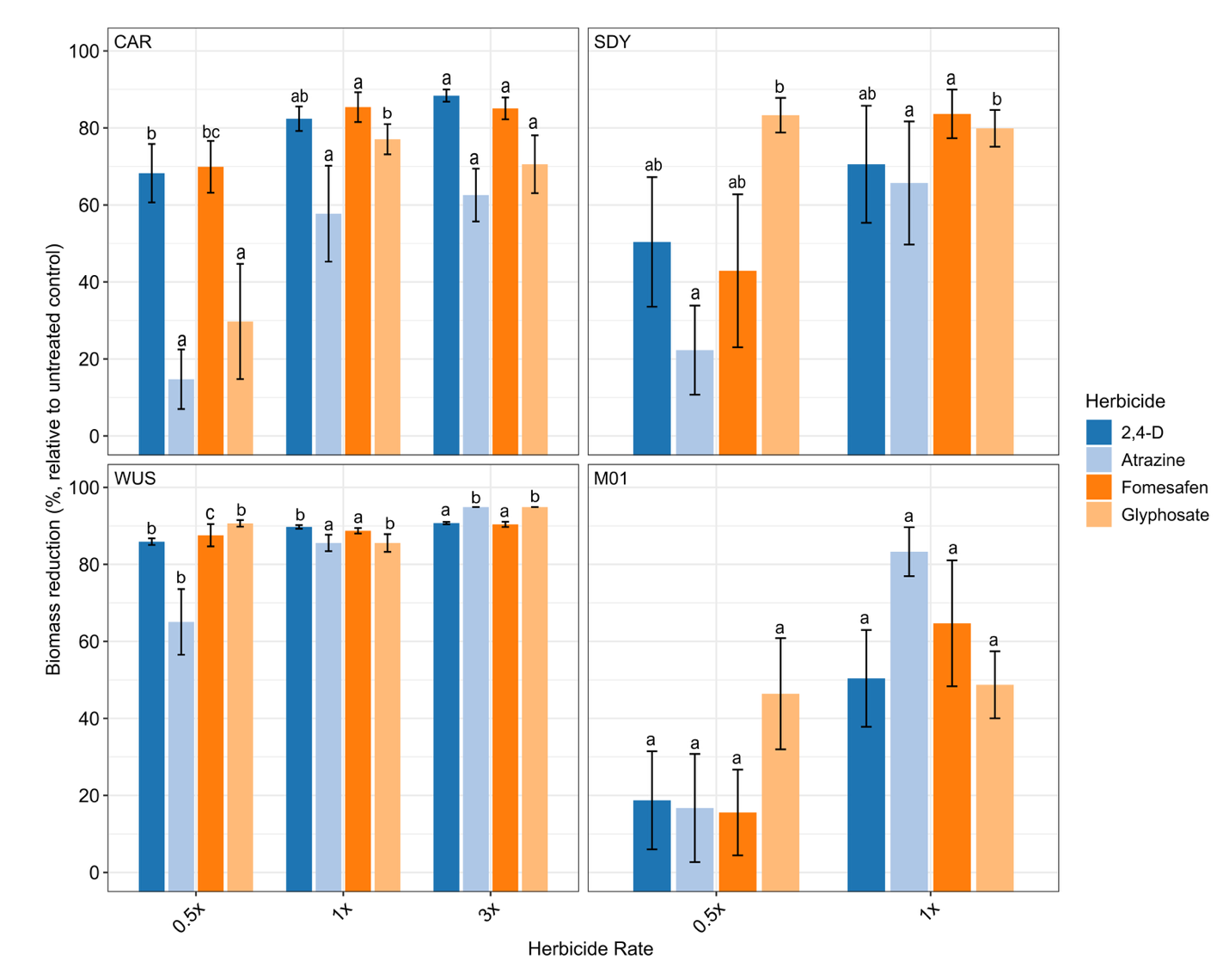

**Supplementary Figure S1.** Biomass reduction relative to the nontreated control (%) of glufosinate-ammonium-resistant (CAR, SDY, and M01) and susceptible (WUS) Amaranthus tuberculatus populations at 21 days after treatment with herbicides representing different sites of action, applied at 0.5×, 1×, and 3× of the field-recommended rate. Bars sharing the same letter are not significantly different according to Tukey’s HSD test (*P* ≤ 0.05).

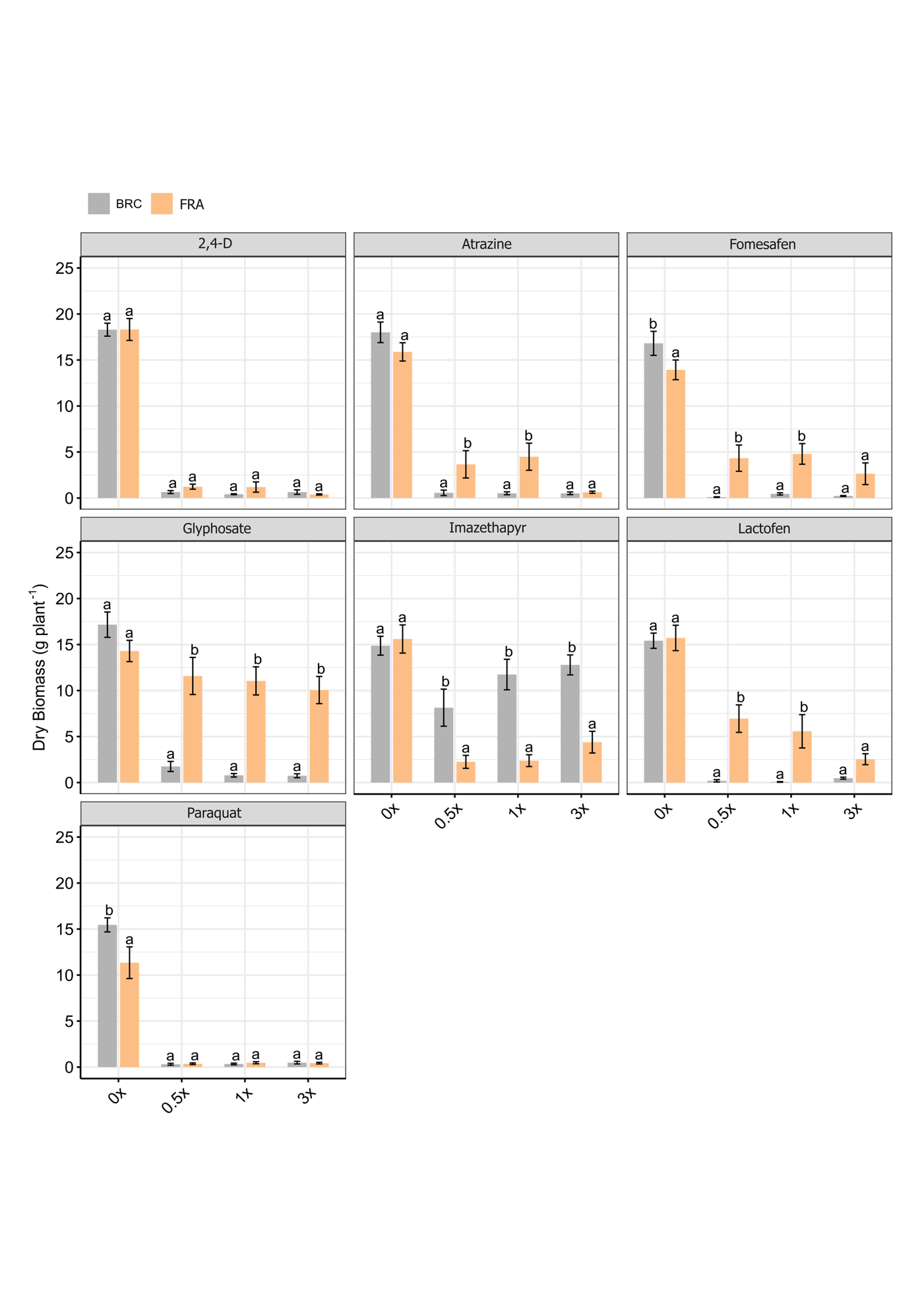

**Supplementary Figure S2.** Dry biomass of glufosinate-ammonium-resistant (FRA) and susceptible (BRC) Amaranthus tuberculatus populations at 21 days after treatment with herbicides representing different sites of action, applied at 0×, 0.5×, 1×, and 3× of the field-recommended rate. Bars sharing the same letter are not significantly different according to Tukey’s HSD test (*P* ≤ 0.05).

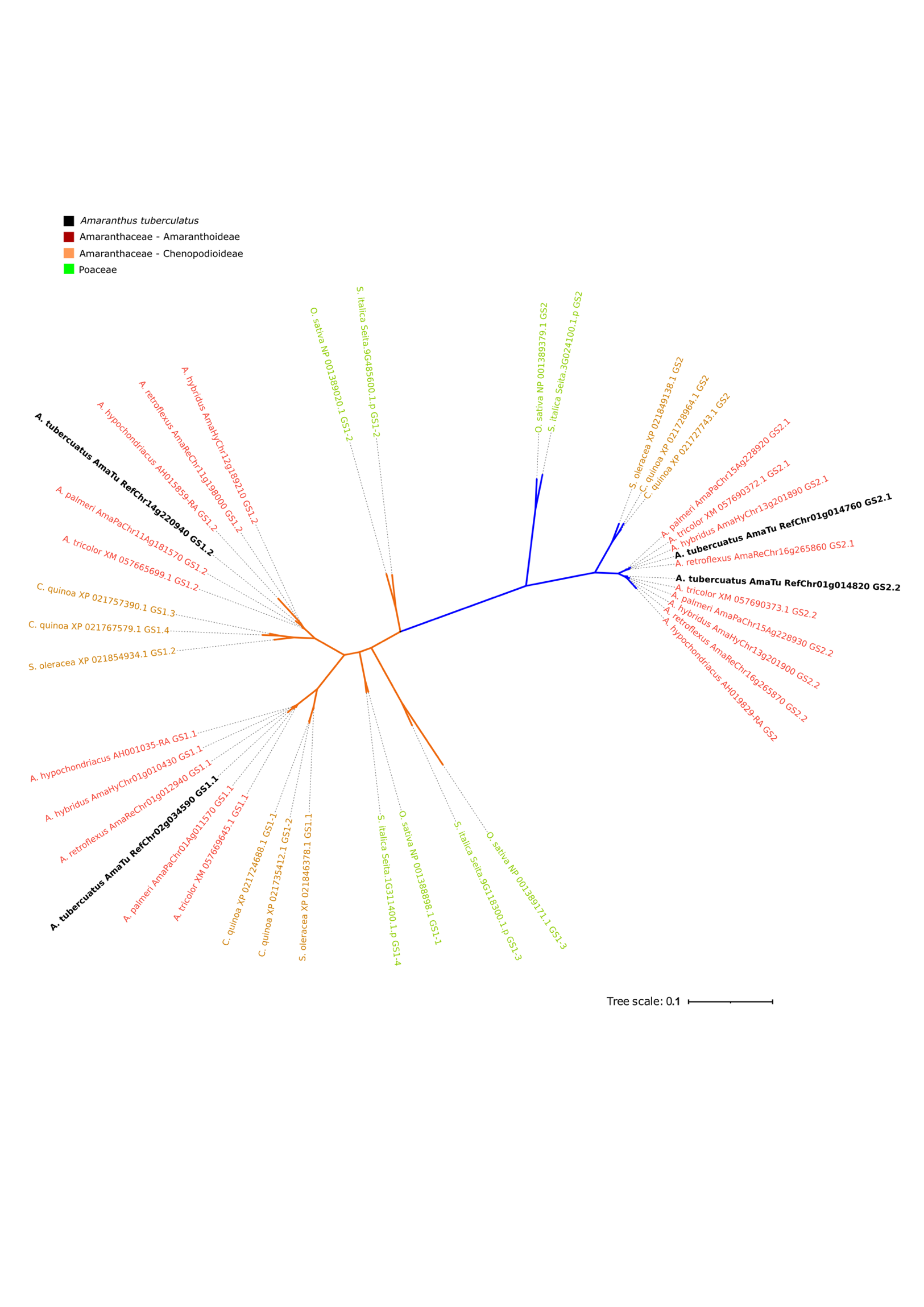

**Supplementary Figure S3.** Phylogenetic relationships among glutamine synthetase isoforms from 10 plant species. Protein sequences were aligned in MEGA, and the resulting phylogenetic tree was visualized using iTOL. Protein identifiers for genes obtained from NCBI and used in phylogenetic analyses are listed in Supplementary Table S2.

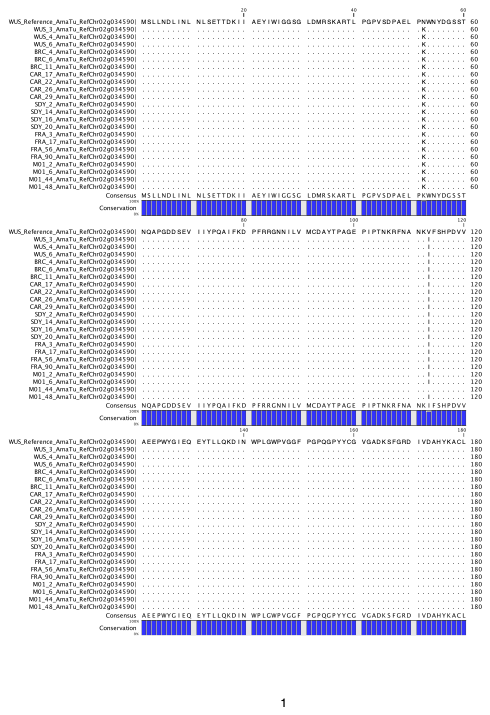

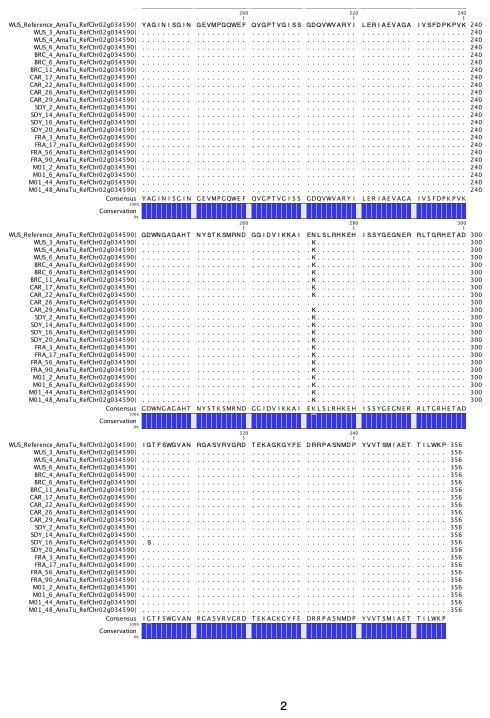

**Supplementary Figure S4.** Multiple sequence alignment of glutamine synthetase (GS1.1) proteins from individual plants of glufosinate-ammonium-resistant (CAR, SDY, FRA, and M01) and susceptible (WUS and BRC) Amaranthus tuberculatus populations. Dots indicate amino acid identity with the reference sequence, and missense mutations are highlighted in yellow. The multiple sequence alignment was generated using CLC Sequence Viewer v8.0.

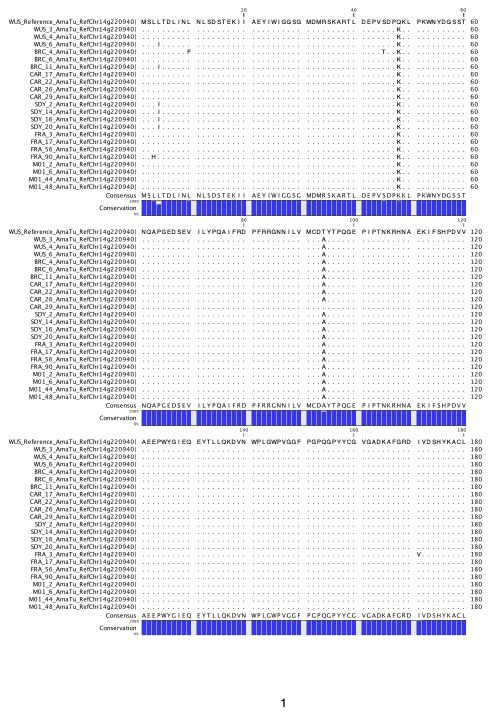

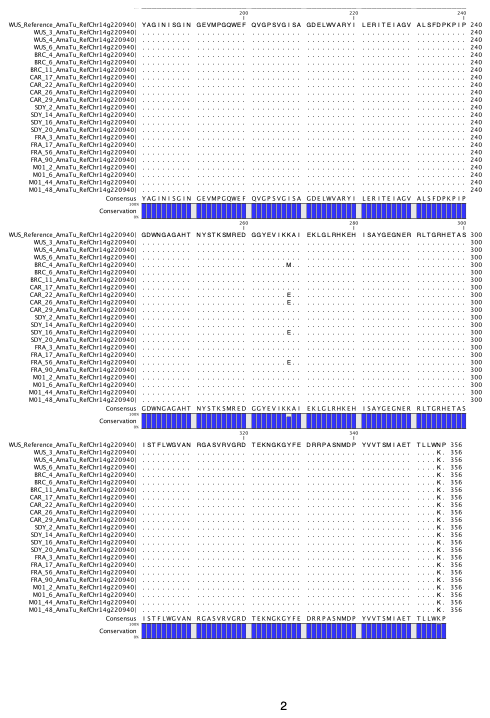

**Supplementary Figure S5.** Multiple sequence alignment of glutamine synthetase (GS1.2) proteins from individual plants of glufosinate-ammonium-resistant (CAR, SDY, FRA, and M01) and susceptible (WUS and BRC) Amaranthus tuberculatus populations. Dots indicate amino acid identity with the reference sequence, and missense mutations are highlighted in yellow. The multiple sequence alignment was generated using CLC Sequence Viewer v8.0.

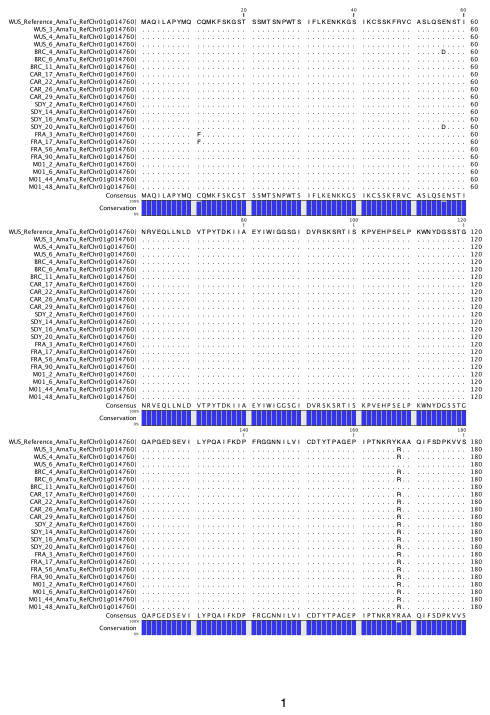

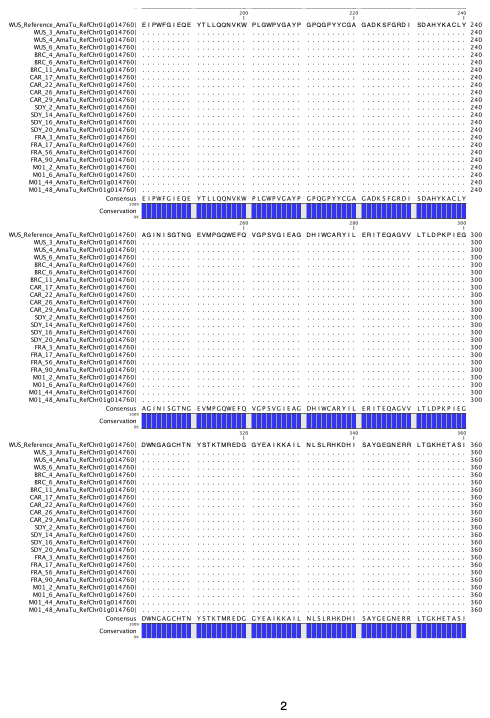

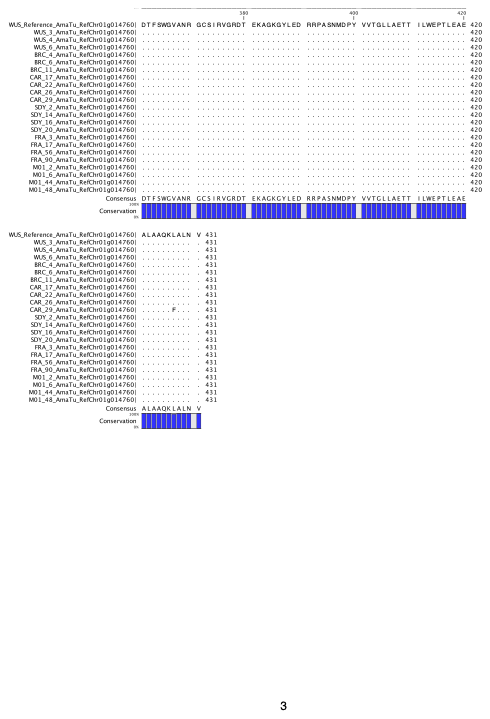

**Supplementary Figure S6.** Multiple sequence alignment of glutamine synthetase (GS2.1) proteins from individual plants of glufosinate-ammonium-resistant (CAR, SDY, FRA, and M01) and susceptible (WUS and BRC) Amaranthus tuberculatus populations. Dots indicate amino acid identity with the reference sequence, and missense mutations are highlighted in yellow. The multiple sequence alignment was generated using CLC Sequence Viewer v8.0.

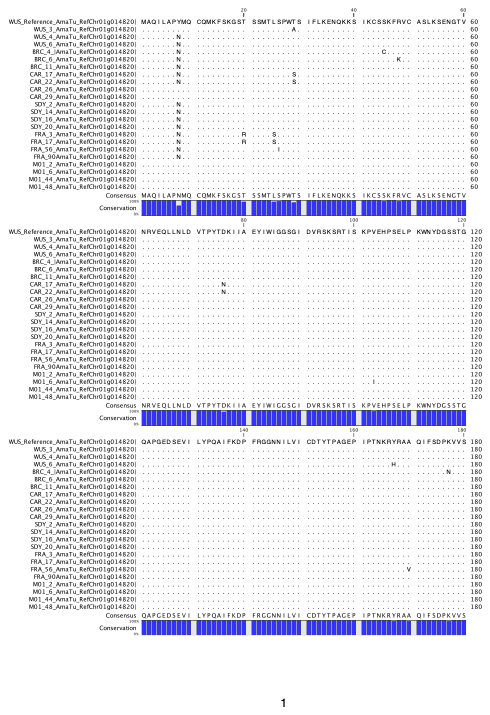

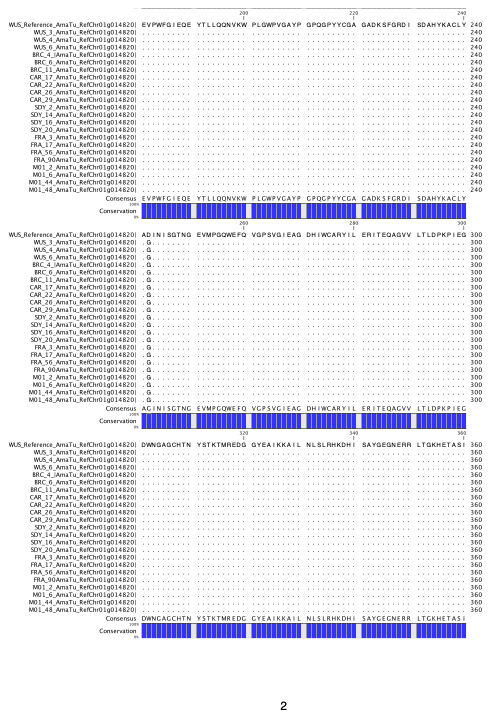

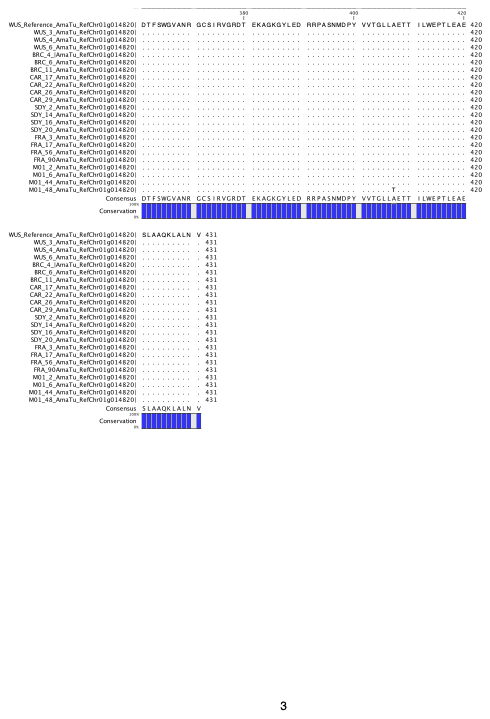

**Supplementary Figure S7.** Multiple sequence alignment of glutamine synthetase (GS2.2) proteins from individual plants of glufosinate-ammonium-resistant (CAR, SDY, FRA, and M01) and susceptible (WUS and BRC) Amaranthus tuberculatus populations. Dots indicate amino acid identity with the reference sequence, and missense mutations are highlighted in yellow. The multiple sequence alignment was generated using CLC Sequence Viewer v8.0.
